# Link between aging and atheroprotection in *Mif*-deficient atherosclerotic mice

**DOI:** 10.1101/2021.12.14.471281

**Authors:** Christine Krammer, Bishan Yang, Sabrina Reichl, Verena Bolini, Corinna Schulte, Heidi Noels, Omar El Bounkari, Aphrodite Kapurniotu, Christian Weber, Sarajo Mohanta, Jürgen Bernhagen

## Abstract

Atherosclerosis is a lipid-triggered chronic inflammatory condition of our arteries and the main underlying pathology of myocardial infarction and stroke. Pathogenesis is age-dependent, but the mechanistic links between disease progression, age, and atherogenic cytokines and chemokines are incompletely understood. Here, we studied the chemokine-like inflammatory cytokine macrophage migration inhibitory factor (MIF) in atherogenic *Apoe*^−*/*−^ mice across different stages of aging and cholesterol-rich high-fat diet (HFD). MIF promotes atherosclerosis by mediating atherogenic monocyte and T-cell recruitment, amplifying lesional inflammation, and suppressing atheroprotective B-cell responses. However, age-related links between atherogenesis and MIF and its role in advanced atherosclerosis in aged mice have not been systematically explored. We compared effects of global *Mif*-gene deficiency in 30-, 42-, and 48-week-old *Apoe*^−*/*−^ mice on HFD for 24, 36, or 42 weeks, respectively, and in 52-week-old mice on a 6-week HFD. While a regio-specific atheroprotective phenotype of *Mif-*deficiency was observed in the 30/24-week-old group, atheroprotection was not detected in the 48/42- and 52/6-week-old groups, suggesting that atheroprotection afforded by global *Mif*-gene deletion differs across aging stages and atherogenic diet duration. We identify a combination of mechanisms that could explain this phenotype: i) *Mif*-deficiency promotes lesional Trem2^+^ macrophage numbers in younger but not aged mice; ii) *Mif*-deficiency favors formation of lymphocyte-rich stage-I/II ATLOs in younger mice but ATLO numbers equalize with those in *Apoe*^−*/*−^ controls in the older mice; and iii) plasma anti-oxLDL-IgM antibody levels are decreased in aged *Mif*-deficient mice. Of note, these three markers (Trem2^+^ macrophages, ATLOs, anti-oxLDL-IgM antibodies) have been previously linked to atheroprotection. Together, our study thus suggests that regio-specific atheroprotection due to global *Mif*-gene deficiency in atherogenic *Apoe*^−*/*−^ mice is lost upon advanced aging and identifies mechanisms that could explain this phenotype shift. These observations may have implications for translational MIF- directed strategies.

## Introduction

Cardiovascular diseases (CVDs) such as myocardial infarction or ischemic stroke are the main underlying causes of death worldwide and incidence risk has steadily increased over the last decades. Disease pathology has been associated with multiple co-morbidities and risk factors such as hypertension, type 2 diabetes (T2D), or metabolic syndrome, which are further impaired by diet and lifestyle changes and increasingly manifest in an aging population (1, 2). Atherosclerosis is a lipid-triggered chronic inflammatory condition of the medium and large arterial vasculature and has been recognized as the main underlying pathology of CVDs. Atherosclerosis may get initiated early in life and progressively develops over years and decades, and is typically clinically symptomatic in later adulthood. Initially triggered by endothelial dysfunction and oxidized low density lipoprotein (oxLDL) uptake, lipid and inflammatory cell deposits in the arterial vessel wall lead to the formation of atheromatous lesions. Lesion formation involves oxLDL-mediated foam cell formation and abundant leukocyte infiltration, a process orchestrated by selectins and adhesion molecules upregulated on the inflamed endothelium as well as by endothelial-deposited chemokines and their receptors and integrins expressed on infiltrating inflammatory cells. Cytokines and chemokines also amplify vascular inflammation through a variety of pathways. Together, this results in increased intima-media thickening, wall remodeling, and limited arterial blood flow, but also eventually necrotic core formation, plaque destabilization, and thrombus formation, consequences which may result in severe adverse clinical outcomes such as myocardial infarction or stroke (3–6).

In addition to classical atherogenic chemokines such as CCL2 or CXCL1/8, atypical chemokines (ACKs) have emerged as players in the atherogenic inflammatory cascade. Atypical chemokines have chemokine-like properties and engage in high-affinity interactions with classical chemokine receptors, but lack classifying structural features of *bona fide* chemokines such as an N-terminal CC- or CXC-motif (7). Macrophage migration inhibitory factor (MIF) is an evolutionarily conserved pleiotropic inflammatory cytokine and prototypical ACK with a potent proatherogenic activity spectrum. MIF is overexpressed in human carotid artery plaques (8) and circulating MIF levels have been associated with coronary artery disease (CAD), suggesting a critical role for MIF in CVDs (9–11). MIF is broadly expressed, but its secretion mainly occurs from immune cells such as monocytes and T cells, as well as endothelial cells, smooth muscle cells (SMCs), and platelets (7, 12). MIF expression is upregulated by various atherogenic stimuli such as inflammatory cytokines, oxLDL, or hypoxia (9, 13). A polymorphism in the CATT microsatellite repeat of the MIF promotor region is associated with higher MIF expression levels in several human inflammatory and autoimmune diseases and an increased susceptibility for carotid artery atherosclerosis (CAA) (14–17). MIF signals through its cognate receptor CD74/invariant chain, but its proatherogenic activities are mainly mediated by non-cognate signaling through the CXC chemokine receptors CXCR2 and CXCR4. This drives the atherogenic recruitment of monocytes, neutrophils, T cells, and platelets (13, 17, 18), and promotes foam cell formation, vascular chemokine and adhesion molecule expression, and arterial wall remodeling (8, 19, 20).

There is a wealth of evidence from *in vivo* models, suggesting a causal role of MIF in atherosclerosis. *Mif*-gene deletion in hyperlipidemic *Ldl receptor*-deficient (*Ldlr*^−*/*−^) mice (18, 21) or antibody-mediated blockade of MIF in *Apolipoprotein e*-deficient (*Apoe*^−*/*−^) mice on a Western-type high-fat diet (HFD) (20, 22) lead to decreased luminal monocyte adhesion, lower plaque macrophage counts, attenuated lesion formation, and increased plaque stability. More recently, Schmitz et *al.* (23) investigated *Mif*-gene deficiency in HFD-fed *Apoe*^−*/*−^ mice and revealed a site-specific atheroprotective phenotype in the brachiocephalic artery (BCA) and abdominal aorta, whereas other regions of the vascular bed were not affected. Moreover, this study provided initial hints about a re-localization of adventitial B cells into cluster-like structures in *Mif*-deficient *Apoe*^−*/*−^ mice but not *Mif*-expressing *Apoe*^−*/*−^ control mice.

Atherosclerosis is an age-dependent pathology driven by chronic inflammation, which becomes clinically symptomatic primarily in advanced stages in aged patients (24). In fact, the tight causal connection between chronic inflammation and aging in CVDs has also been coined “inflamm-aging” (25). In the *Apoe*^−*/*−^ mouse model of atherosclerosis, the development of advanced atheromatous plaques can be recapitulated within 12-24 weeks of HFD (26), but this and similar models do not fully mirror the advanced and late stage vessel phenotypes and clinical manifestations observed in humans, and atherogenic mice at an advanced age of >32 weeks, which would be equivalent to an age span of >50 years in humans (https://www.jax.org/research-and-faculty/research-labs/the-harrison-lab/gerontology/life-span-as-a-biomarker; (27)), have been investigated rather rarely. Furthermore, while studies on the impact of aging in atherosclerosis have identified contributing factors such as dysregulated cytokines and chemokines or vascular mitochondrial dysfunction, the underlying regulatory mechanisms and the details of their interplay with disease progression have remained incompletely understood. Interestingly, artery (or adventitial) tertiary lymphoid organs (ATLOs) have emerged as atherosclerosis-associated lymphoid clusters with disease-modulating activity (28, 29). ATLOs are peri-adventitial B- and T-lymphocyte-rich cell clusters with a lymphoid organ-like structure, which, in the *Apoe*^−*/*−^ mouse model, are typically observed to develop in 52-78-week-old mice (28). They have been suggested to especially control the B-cell response in aged atherosclerotic mice (30). ATLOs have also been detected in patients with CAD, but not much is currently known about their formation and disease mechanism (31). Overall, these observations highlight the important link between aging and the dysregulation of the immune and inflammatory response in atherosclerosis. However, the effects of MIF activity have not been studied in the context of aging.

Here we sought to characterize age-related effects of global *Mif*-gene deletion in advanced stages of atherosclerosis. This was studied by comparing lesion formation, leukocytes profiles, and systemic inflammation between *Mif*^−*/*−^ *Apoe*^−*/*−^ and *Apoe*^−*/*−^ mice that were on HFD for 24, 36 and 42 weeks, corresponding to an age of 30, 42, and 48 weeks, respectively, as well as in 52-week-old mice on HFD for 6 weeks. These age/HFD groups are hereinafter termed the 30/24-, 42/36-, 48/42-, and 52/6-week-groups. The impact of B cells was investigated by an in-depth analysis of peri-adventitial cell clusters and antibody profiles directed against oxLDL. We confirm the regio-specific atheroprotective effect of *Mif*-deficiency in BCA and abdominal aorta in *Apoe*^−*/*−^ mice in the 30/24-week group. Notably, the site-specific atheroprotective effect was reduced in the 48/42-week group as well as in the 52/6-week group. This was accompanied by MIF- and age-dependent changes in plaque immune cells, peri-adventitial cell clusters, and atheroprotective natural IgM antibodies. Together, our study for the first time reveals age-dependent functions of the atypical chemokine MIF in atherosclerotic pathology and suggests that a combination of mechanisms could account for reduced atheroprotection in *Mif*-deleted aged atherogenic *Apoe*^−*/*−^ mice.

## Materials and methods

### Chemicals, buffers, and miscellaneous reagents

Miscellaneous reagents were purchased from Sigma Aldrich/Merck (Darmstadt, Germany), VWR International GmbH (Darmstadt, Germany), Carl Roth GmbH (Karlsruhe, Germany), and Ratiopharm (Ulm, Germany) and were of the highest purity degree available.

### Mice

Atherosclerosis-prone *Apolipoprotein e*-deficient (*Apoe*^−*/*−^) mice and *Apoe^−/–^Mif^−/–^* mice were in the C57BL/6-J background. *Apoe*^−*/*−^ mice were initially obtained from Charles River Laboratories (Sulzfeld, Germany) and were backcrossed within the animal facility of the Center for Stroke and Dementia Research (CSD). Global *Mif* gene-deficient mice (*Mif^−/–^*) were generated by Dr. Fingerle-Rowson and Prof. Richard Bucala and have been reported on before (23, 32). For the generation of *Mif^−/–^Apoe^−/–^* mice, *Mif^−/–^* mice were backcrossed with *Apoe^−/–^* mice for at least 10 generations. Mice were housed and bred under standardized and specific pathogen-free conditions in the animal facility of the CSD in Munich, with free access to food and water. Animal experiments were approved by the local authorities (animal ethics approval ROB-55.2-2532.Vet_02-18-040) and were performed according to the German animal protection law. Animals were sacrificed under anesthesia with a mixture of midazolam (5 mg/ml), medetomidine and fentanyl (MMF).

### Western-type high-fat diet (HFD)

Female animals received a regular chow diet (SNIFF Spezialdiäten GmbH, Soest, Germany) until 6 weeks of age and then were subjected to a high-cholesterol (‘Western type’) diet (HFD) containing 0.2% cholesterol and 21.2% total fat (TD88137, SNIFF Spezialdiäten GmbH, Soest, Germany) for an additional 24, 36, or 42 weeks. Mice aged for 52 weeks received the HFD only in the last 6 weeks before the end of the experiment.

### Isolation of blood, organs, and vessels

Blood was collected by cardiac puncture into ethylene diamine tetra-acetic acid (EDTA)- containing tubes (Sarstedt, Nümbrecht, Germany) and hematologic parameters were subsequently analyzed using the Scil Vet aBCA Plus+ Blood Analyzer (Scil Animal Car Company GmbH, Viernheim, Germany). For plasma preparation, the blood was centrifuged at 400 x g for 15 min at 4°C. Plasma samples were immediately frozen in liquid nitrogen and stored at -80°C. The circulation was rinsed with 15 ml of perfusion solution (100 U/ml heparin, 10 mM EDTA in phosphate-buffered saline (PBS), pH 7.4) followed by 15 ml of PBS.

Single cell suspensions were generated from spleen, lymph nodes (LNs) and bone marrow (BM) of femur and tibia by filtering the cells through a 40 µm cell strainer (Corning, Sigma Aldrich/Merck). After red blood cell (RBC) lysis using RBC lysis buffer (BioLegend, Koblenz, Germany), cells were washed in PBS and used for subsequent analysis. Different parts of the vascular bed, including the brachiocephalic artery (BCA) and the aortic root, were isolated and embedded in Tissue-Tek® O.C.T.™ compound (Sakura Finetek, Staufen, Germany). The embedded tissues were stored at -80°C until preparation of 5 µm thick serial cryosections. The whole aorta was excised, pinned onto a slide and fixed overnight in 1% paraformaldehyde (PFA) at 4°C. The next day the aorta was removed from surrounding fat and *en face* preparations were generated. The adventitia was removed and the aorta pinned with the endothelium pointing upwards. The *en face* prepared vessel was again fixed in 4% PFA overnight and then applied to subsequent analyses.

### Flow cytometric analysis

Flow cytometric analysis was performed to analyze the immune cell content (CD8^+^ T cells: CD45^+^CD3e^+^CD8a^+^, CD4^+^ T cells: CD45^+^CD3e^+^CD4^+^, monocytes: CD45^+^CD11b^+^CD115^+^, neutrophils: CD45^+^CD11b^+^Ly6G^+^, B cells: CD45^+^CD19^+^) of different organs including BM, spleen, LN, and blood using a BD FACSVerse^TM^ instrument with a 3 laser, 8 color (4-2-2) configuration (BD Bioscience, Heidelberg, Germany). The system automatically adjusts spillover values for the compensation of standard fluorochromes. Cells were stained with fluorescently-labeled antibodies directed against cell-specific surface marker for 30 min one ice in the dark (anti-mouse CD45-APC/Cy7, #103116, BioLegend; anti-mouse CD45-FITC, #FAB114F, R&D Systems, Wiesbaden-Nordenstadt, Germany; anti-mouse CD3e-FITC, #130- 119-758, Miltenyi Biotech, Bergisch Gladbach, Germany; anti-mouse CD4-PE, #130-102-619, Miltenyi Biotech; anti-mouse CD8a-PE-Cy7, #25-4321-82, ThermoFisher Scientific, Darmstadt, Germany; anti-mouse CD19-PerCP-Cy5.5, #115534, BioLegend; anti-mouse CD11b-FITC, #130-081-201, Miltenyi Biotech; anti-mouse CD115-PE, #12-1152-82, ThermoFisher Scientific; anti-mouse Ly6G, # 45-5931-80, ThermoFisher Scientific). The antibodies were diluted 1:100 in FACS staining buffer (0.5% bovine serum albumin (BSA)/PBS, pH 7.4). Afterwards, the cells were washed with FACS buffer and centrifuged for 5 min at 300 x g and 4°C and flow cytometric analysis was performed. For control, cells were stained with respective isotype control antibodies purchased from BioLegend or R&D Systems (rat IgG2b,κ-APC/Cy7, rat IgG2b,κ-FITC, rat IgG2a,κ-PE, rat IgG2b,κ-PerCP/Cy5.5, rat IgG2a,κ- PerCP/Cy5.5, rat IgG2b,κ-PE, rat IgG2a,κ-PE/Cy7) or REA control antibodies (REA Control (I)-FITC, Miltenyi Biotech). Results were analyzed by FlowJo software (Tree star, USA) and presented as dot plots at a logarithmic scale.

### Plasma anti-oxLDL antibody titer

Plasma anti-oxLDL antibody levels were measured in Nunc MaxiSorp^TM^ 96-well plates (ThermoFisher Scientific) coated with 1 µg/ml oxLDL (ThermoFisher Scientific) diluted in carbonate buffer (34.8 mM NaHCO3, 15 mM Na2CO3 in double-distilled (dd)H2O). After coating for 1 h at 37°C, plates were washed three times with 100 µl of washing/blocking solution (2% BSA in PBS) and blocked for 1 h at 37°C. Next, 50 µl plasma diluted 1:100 in washing/blocking solution was added per well and incubated for 1 h at 37°C. The plates were washed again and 50 µl of horseradish-peroxidase (HRP)-labelled antibodies against IgG (# ab6789, Abcam, Berlin, Germany) and IgM (#62-6820, ThermoFisher Scientific) diluted 1:500 in washing-/blocking solution were applied and incubated for 1 h at 37°C. Following additional washing steps, the assay was developed by adding 100 µl Pierce^TM^ 3,3’,5,5’-tetramethylbenzidine (TMB) substrate solution (ThermoFisher Scientific) for 5 min and the reaction was stopped by applying 50 µl stop solution (0.5 M H2SO4 in ddH2O). Measurements were performed using an EnSpire Multimode Plate Reader (PerkinElmer, Hamburg, Germany) at 450 nm.

### Plaque morphometry, immunofluorescence, *en face* staining of aorta, and plaque lipids

#### Immunofluorescence of brachiocephalic artery

Immunohistochemistry was performed on 5 µm thick cryosections of the BCA. The sections were fixed with pre-cooled acetone at 4°C for 6 min and air-dried at room temperature (RT) for 30 min, following rehydration in PBS for 10 min. Next, the sections were blocked for 30 min with blocking solution (5% donkey/goat serum, 1% BSA in PBS, pH 7.4). Next, the sections were incubated with the primary antibodies diluted in blocking solution at 4°C overnight. The following primary antibodies were applied: rat anti-mouse CD45R/B220 (1:100, #557390, BD Biosciences), rat anti-mouse CD68 (1:100, #MCA1957GA, Bio-Rad, Puchheim, Germany), hamster anti-mouse CD3e (1:100, #553058, BD Biosciences), mouse anti-human/mouse SMA-Cy3 (1:200, #C6198, Sigma Aldrich), rat anti-mouse CD35 (1:100, #558768, BD Biosciences), rat anti-mouse PNAd (1:50, #553863, BD Biosciences), rabbit anti-mouse collagen IV (1:500, #2150-1470, Bio-Rad), rat anti-mouse ER-TR7 (1:500, #ab51824, Abcam), rabbit anti-mouse Lyve1 (1:500, #DP35135P, Acris Antibody GmbH, Herford, Germany) and sheep anti-mouse Trem2 (1:50, #AF1729, R&D Systems), and rabbit anti-mouse Ki67 (1:200, #9129T, Cell Signaling). Afterwards, sections were washed in PBS and incubated for 1 h at RT with the respective secondary antibodies diluted in blocking solution. The following secondary antibodies were used: donkey anti-rat-Cy5 (1:300, Jackson ImmunoResearch, Hamburg, Germany, #712-175-153), goat anti-rat-AF488 (1:500, #A-11006, ThermoFisher Scientific), goat anti-hamster-Cy3 (1:300, #127-165-160, Jackson ImmunoResearch), goat anti-rat(IgM)-Cy5 (1:500, #A21247, ThermoFisher Scientific), donkey anti-rabbit-Cy3 (1:500, #711-165-152, Jackson ImmunoResearch), donkey anti-rat-Cy3 (1:300, #712-166-153, Jackson ImmunoResearch), donkey anti-sheep-AF555 (1:500, ab150178, Abcam), and donkey anti-sheep-AF647 (1:500, ab150179, Abcam), and goat anti-rabbit-AF647 (1:300, #A21245, ThermoFisher Scientific). DAPI was used as nuclear counterstain. Afterwards, sections were washed and mounted with Fluoromount^TM^ aqueous mounting medium (Sigma-Aldrich/Merck). Images were recorded with a DMi8 fluorescent microscope or a TCS SP5II MP confocal microscope (Leica Microsystems, Wetzlar, Germany) and analyses were performed using ImageJ software.

#### Oil-Red-O staining of aortic root

For the quantification of lesion formation in the aortic root, Oil-Red-O (ORO) stainings were performed on 5 µm thick cryosections. First, the slides were air-dried for 5 min following rehydration in PBS for 2 min. Next, lipids were stained with ORO solution (0.5% in propylene glycol, Sigma Aldrich/Merck) at 37°C for 45 min. The slides were shortly washed with running tap water and nuclear counterstain was performed using hematoxylin. The slides were air-dried and mounted with Kaiser’s glycerin gelatine mounting media (Carl Roth, Karlsruhe, Germany). Images were taken with the DMi8 fluorescent microscope and lesion quantification performed using ImageJ software. Per mouse, 12 serial sections with a distance of 50 µm between each other, were stained, and mean values calculated.

#### Oil-Red-O (en face) staining of aorta

Oil-Red-O stainings of the aorta including aortic arch as well as abdominal and thoracic aorta were performed after *en face* preparation and fixation in 4% PFA. The tissue was stained for 30 min in ORO solution (0.5% in isopropanol, Sigma Aldrich/Merck) at RT. Next, aortas were washed in 60% isopropanol until unspecific ORO-derived staining was removed from the endothelium. The aortas were rinsed briefly with running tap water before mounting in Kaiser’s glycerin gelatine mounting media. Tilescan images were acquired with the DMi8 fluorescent microscope. The lesions were quantified via FlowJo software and depicted as percentage of total aortic surface.

#### Hematoxylin-eosin staining of brachiocephalic artery

Hematoxylin-eosin (H&E) staining was performed for lesion quantification of the BCA. The slides were air-dried at RT for 30 min and rehydrated with PBS for 2 min. Next, nuclei were stained with Mayer’s hematoxylin solution (Sigma Aldrich/Merck) for 15 min at RT. Then, the slides were washed in running tap water for 10 min before staining with Eosin G-solution for 10 sec and dehydration by two changes of 95% ethanol, 100% ethanol and xylene (2 min each). After air-drying, the slides were mounted using a VECTASHIELD® antifade mounting medium (Vector Laboratories). Images were taken with a Leica DMi8 or a Leica DM6 B fluorescent microscope. The lesion size was quantified using ImageJ software and is depicted as ratio of total inner vessel area. Per mouse, 12 sections with a distance of 50 µm between them were stained and the mean values calculated.

### MIF mRNA expression analysis

#### Trizol-based RNA isolation

For the analysis of MIF mRNA levels, RNA was extracted from frozen tissues using a TRIzol^TM^- based protocol. First, tissue was homogenized in 1 ml of TRIzol^TM^ reagent (ThermoFisher Scientific) using stainless steel beads (5 mm mean diameter) and a TissueLyser LT adapter (Qiagen, Hilden, Germany) for 5 min at 50 Hz. Afterwards, tissue lysates were incubated for 5 min to permit complete dissociation of nucleoprotein complexes. After adding 200 µl of chloroform and incubation for 3 min, the solution was centrifugation for 15 min at 12000 x g and 4°C. The RNA-containing fraction was mixed with 250 µl isopropanol and incubated for 10 min. Next, the samples were centrifuged at 10000 x g and 4°C for 10 min and the RNA pellets resuspended in 75% ethanol. After centrifugation for 5 min at 7500 x g and 4°C, the RNA pellets were air-dried and the RNA dissolved in nuclease-free water. The RNA concentration was measured using a NanoDrop^TM^ One UV/Vis spectrophotometer (ThermoFisher Scientific).

### Reverse transcription and quantitative real-time PCR (RT-qPCR)

To transcribe the purified RNA into single stranded cDNA, the First Strand cDNA synthesis kit (Thermo Fisher Scientific) was used according to the manufactureŕs protocol. Briefly, a reaction mix containing 1 µl RNase inhibitor, 2 µl dNTP mix, 2 µl reverse transcriptase,1 µl Oligo(dT)18 primers and 4 µl 5 x reaction buffer was prepared and mixed with 1 µg of isolated RNA. The reverse transcription was performed in a Biometra thermocycler (Analytik Jena AG, Jena, Germany) with following incubation settings: 1 h at 37°C, 5 min at 70°C, cooling down to 4°C. RT-qPCR was performed using ORA^TM^ SEE qPCR Green ROX H Mix (HighQu, Kraichtal, Germany) and specific mouse primer pairs (Eurofins, Ebersberg, Germany). The following primers were used: *MIF forward: ACA GCA TCG GCA AGA TCG* and *MIF reverse: AGG CCA CAC AGC AGC TTA C; actin forward: GGA GGG GGT TGA GGT GTT* and *actin reverse: GTG TGC ACT TTT ATT GGT CTC AA*. PCR reactions were run in a Rotorgene Q (Qiagen). Relative mRNA levels were calculated using the ΔΔCt method with *actin* as a housekeeping gene.

### Lipid analysis

#### Cholesterol fluorometric assay

Cholesterol levels in plasma were measured using Cayman’s Cholesterol Fluorometric Assay kit (Cayman Chemicals/ Biomol GmbH, Hamburg, Germany) according to the manufacturer’s instructions. Briefly, plasma was diluted 1:2000 in cholesterol assay buffer and 50 µl of the samples were added to the wells of a 96-well plate. After initiation the reaction with 50 µl of assay cocktail (4.745 ml cholesterol assay buffer, 150 µl cholesterol detector, 50 µl cholesterol assay HRP and 5 µl cholesterol esterase), the plates were incubated for 30 min at 37°C in the dark. Afterwards the fluorescence was measured at an excitation wavelength of 535 nm and an emission wavelength of 590 nm. Samples were measured in duplicates using an EnSpire Multimode Plate Reader (PerkinElmer). Diluted cholesterol standards with final concentrations between 2 and 20 µM were used for the quantification of cholesterol levels in the plasma.

#### Triglyceride colorimetric assay

Triglyceride levels in plasma were measured using Cayman’s Triglyceride Colorimetric Assay kit (Cayman Chemicals/ Biomol GmbH) according to the manufacturer’s instruction. Briefly,10 µl of undiluted plasma was mixed with 100 µl of pre-diluted enzyme mixture in a 96-well plate and incubated for 15 min at RT. The absorbance was measured at 540 nm using the EnSpire Multimode Plate Reader (PerkinElmer). Diluted triglyceride standards with final concentrations between 3.125 and 200 mg/dl were used for the quantification of triglyceride levels in mouse plasma.

#### Murine MIF ELISA

MIF levels in murine plasma were measured by the Mouse MIF DuoSet® ELISA (R&D Systems) according to the manufacturer’s instruction. Briefly, wells of a 96-well plate were coated with 100 µl of capture antibody at a final concentration of 400 ng/ml overnight at RT. After three washes with washing buffer (0.05% Tween 20 in PBS, pH 7.2), plates were blocked in 300 ml of 1x reagent diluent concentrate 2 (R&D Systems) for 1 h at RT. After additional washing, 100 µl of the appropriately diluted plasma sample or standard in 1x reagent diluent concentrate 2 were added into each well. Standards were prepared using the recombinant mouse MIF standard supplied with the kit using 2-fold serial dilutions. Standard concentrations ranged from 125 pg/ml to 2000 pg/ml. After incubation with samples/standards for 2 h at RT, washing was repeated as described, and 100 µl of detection antibody at a final concentration of 200 ng/ml were added for 2 h at RT. Plates were washed three times, 100 µl of Streptavidin-HRP (1:200 dilution) added, and incubated for 20 min at RT, followed by washing and incubation with 100 µl of TMB substrate solution (ThermoFisher Scientific) for another 20 min at RT. The reaction was stopped by adding 50 µl of stop solution (2N H2SO4) per well. Optical density was determined at 450 nm using an EnSpire Multimode Plate Reader (PerkinElmer).

Plasma dilutions of 1:50 and 1:100 were found to be optimal for best signal/noise ratios and mean values were used for analysis.

### Statistics

All statistical analyses were performed using GraphPad Prism 6.0 or 7.0 (GraphPad Software Inc.). Data are represented as means ± SD. After testing for normality using the *D’Agostino-* Pearson test, data were analyzed by two-tailed Student’s T-test (parametric) or by Mann-Whitney test (non-parametric) as appropriate. In case of comparing more than two groups, two-way ANOVA with Sidak’s multiple comparisons test was applied. *P*<0.05 was considered statistically significant.

## Results

### Site-specific atheroprotection due to global *Mif-*deficiency in *Apoe***^−^***^/^***^−^** mice is reduced upon aging

The role of MIF in atherogenesis has previously been investigated in younger and middle-aged atherogenic *Apoe*^−*/*−^ or *Ldlr*^−*/*−^ mice on HFD for up to 14-26 weeks (18, 20–23, 33). However, age-dependent effects of MIF were not addressed in these studies. Here, we determined the effect of global *Mif*-deficiency in 30-, 42-, and 48-week-old *Apoe*^−*/*−^ mice that received an HFD for 24, 36 and 42 weeks, respectively, as well as in 52-week-old mice that received an HFD in the last 6 weeks, i.e. in 30/24-, 42/36-, 48/42-, and 52/6-week treatment groups.

We first compared the body weights of *Apoe*^−*/*−^*Mif*^−/–^ mice with those of *Apoe*^−*/*−^ controls over the course of aging and HFD. *Mif*-deficiency appeared to slightly affect weight gain across the course of aging. Whereas the body weight of *Apoe*^−*/*−^ control mice increased with age and the duration of the HFD (24 weeks HFD: 35.4±4.8 g; 42 weeks HFD: 38.2±5.5 g), the *Mif*- deficient animals maintained or even lost body weight over the same course (24 weeks HFD: 34.0±3.7 g; 42 weeks HFD: 32.6±3.3 g), but this trend did not reach statistical significance (**Supplementary Table S1**). This observation was paralleled by corresponding differences in lipid levels (**Supplementary Table S2**), and significantly decreased triglyceride levels were detected in the plasma of *Apoe*^−*/*−^*Mif*^−/–^ mice after 42 weeks of HFD compared to *Apoe*^−*/*−^ controls (**Supplementary Table S2**; *Apoe*^−*/*−^*Mif*^−/^: 116.9±28.9 g; *Apoe*^−*/*−^: 211.4±88.2 g; *P*=0.025). Cholesterol levels in the *Apoe*^−*/*−^*Mif*^−/–^ mice were lower both after 24 and 42 weeks of HFD compared to *Apoe*^−*/*−^ controls, but the differences did not reach statistical significance (**Supplementary Table S2**). Also, additional blood parameters such as total white blood cell count (WBC), red blood cells (RBC), hemoglobin (HGB), and hematocrit (HCT) did not differ between mouse strains and age groups (**Supplementary Table S3**).

We next analyzed the atherosclerotic lesions across the vascular tree. Plaque area analysis in 30/24-week group confirmed the intriguing regio-specific atheroprotective effect of *Mif*-gene deletion that had been previously observed in atherogenic mice with an *Apoe*^−*/*−^ background (23). *Apoe*^−*/*−^*Mif*^−/–^ mice exhibited a significantly reduced plaque size compared to *Apoe*^−*/*−^ controls in BCA and abdominal aorta, but not in aortic root, aortic arch, and thoracic aorta (**Figure 1**). Notably, the atheroprotective effect of *Mif*-knockout was lost over the time course of aging and HFD. While attenuation of plaque formation in the *Mif*-knockout genotype was still observed in the BCA and abdominal aorta after 36 weeks of HFD, it was lost in the 48/42-week group (**Figure 1a,b,e,f,i**), with an almost inverted plaque phenotype seen in abdominal aorta of the most aged animals of 48/42 weeks (**Figure 1i**). The loss of the atheroprotective phenotype in *Mif*-deficient mice at an advanced age of 48 weeks was further confirmed in the 52/6-week mouse group. No difference in plaque size was seen in those mice, neither in aortic root, arch, and thoracic aorta, nor in BCA and abdominal aorta (**Supplementary Figure S1a-h**). The similar outcome in the 48/42- and 52/6-week models could be indicative of a predominant role of aging *per se* over the duration of the HFD in an atherogenic *Apoe*^−*/*−^ background.

**Figure 1:**
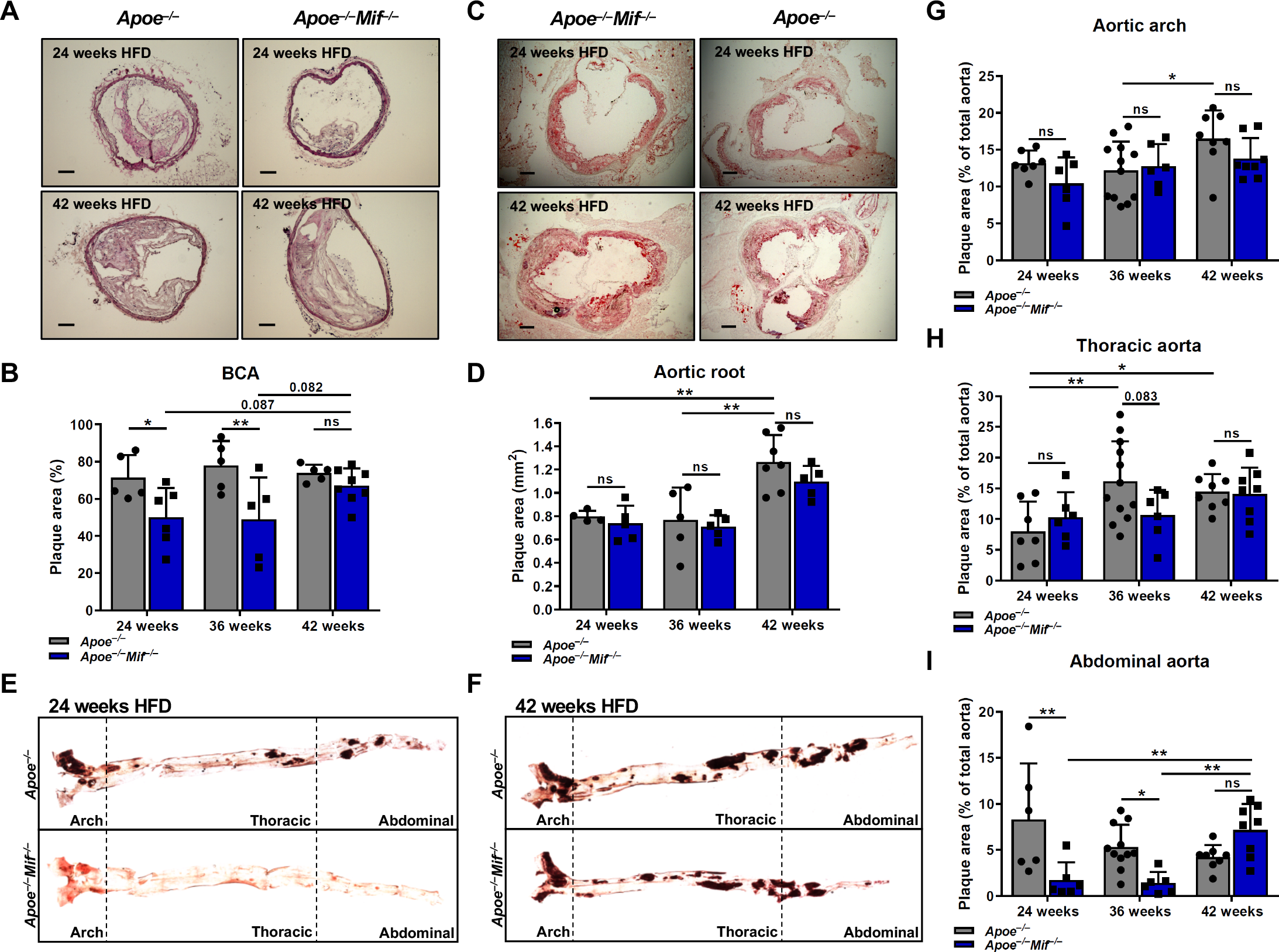
Atheroprotection due to *Mif*-deficiency is lost in highly-aged hyperlipidemic *Apoe^−/–^* mice. Atherosclerotic plaques were quantified in BCA, aortic root, aortic arch, thoracic and abdominal aorta of 30-, 42-, and 48-week-old *Apoe^−/–^Mif^−/–^* mice after 24, 36, and 42 weeks of HFD, respectively, (blue) and compared to corresponding lesions in *Apoe^−/–^* mice (grey). **A-B**) Representative images (**A**) and plaque quantification (**B**) of H&E-stained BCA sections. For each mouse, 12 serial sections with a distance of 50 µm were used for analysis. The mean plaque area is depicted as percentage of the total inner vessel area including the plaque (n=5- 8, results are presented as means ± SD; scale: 100 µm). **C-D**) Representative images (**C**) and quantification (**D**) of ORO-stained sections of the aortic root. Serial sections were obtained as in (**A-B**). The mean plaque area is depicted in mm^2^ (n=4-7, results are shown as means ± SD; scale: 200 µm). **E-F**) Representative images of *en face-*prepared and ORO-stained aortas after 24 (**E**) and 42 (**F**) weeks of HFD. **G-I**) Quantification of plaque area in aorta including aortic arch (**G**), thoracic aorta (**H**) and abdominal aorta (**I**). The plaque area is depicted as percentage of the total aortic surface (n=6-12, results presented as means ± SD). Statistics: two-way ANOVA; *, *P*<0.05; **, *P*<0.01; ns, non-significant; non-significant results with *P* values between 0.05 and 0.1 are given as precise numbers; each data point represents one independent mouse).

MIF expression levels may decrease over the course of aging (34) and this may lead to a relative reduction in the difference of MIF levels between *Apoe*^−*/*−^*Mif*^−/–^ and *Apoe*^−*/*−^ mice, in turn affecting the differences seen in atheroprotection at different age stages. However, quantification of MIF expression levels in liver and spleen by qPCR as well as plasma MIF determinations by MIF ELISA did not reveal differences in MIF levels in *Apoe*^−*/*−^ mice after 24 *versus* 42 weeks of HFD, i.e. in 30- *versus* 48-week-old mice (**Supplementary Figure S2**).

The comparison of plaque phenotypes across age, HFD duration, and vascular bed locations led to additional interesting observations irrespective of the investigation of *Mif*- deficiency. In the BCA of *Apoe*^−*/*−^ mice, plaque burden stagnated at approximately 75% plaque area after 24 weeks of HFD (**Figure 1b**), whereas an aging/HFD duration-dependent increase was observed in aortic root, arch, and thoracic aorta (**Figure 1d,g,h**). A trend towards a decrease in plaque burden was noted in the abdominal part of the aorta (**Figure 1i**).

In conclusion, the atherosclerotic lesion analysis confirmed that *Mif*-deficiency attenuates lesion formation in a regio-specific manner in the relatively younger mice and showed that the atheroprotective effect of global *Mif*-deletion is reduced or lost at an advanced age. Thus, aging seems to be a contributing factor, when considering MIF-mediated effects during atherosclerotic disease progression.

### CD4^+^ T-cell numbers in spleen and blood are changed in an age- and MIF-dependent manner

To begin to explore the mechanism underlying the observed age- and MIF-dependent plaque phenotype, we applied flow cytometry to analyze immune cell populations in spleen, blood, LNs, and BM. Cell numbers were compared for the 24- and 42-week HFD treatment regimens between *Apoe*^−*/*−^ and *Apoe*^−*/*−^*Mif*^−/–^ mice. Interestingly, the most significant changes were seen for T cells. Total splenic T-cell levels were significantly reduced in *Apoe*^−*/*−^ mice over the course of aging, and a sub-population analysis showed that this effect was due to a decrease in CD4^+^ T cells, whereas CD8^+^ T-cell numbers remained unchanged (**Figure 2a**). A similar effect was observed for circulating T cells, although differences did not reach statistical significance, while no changes were noted for T cells isolated from LNs or BM (**Figure 2b-d**).

**Figure 2:**
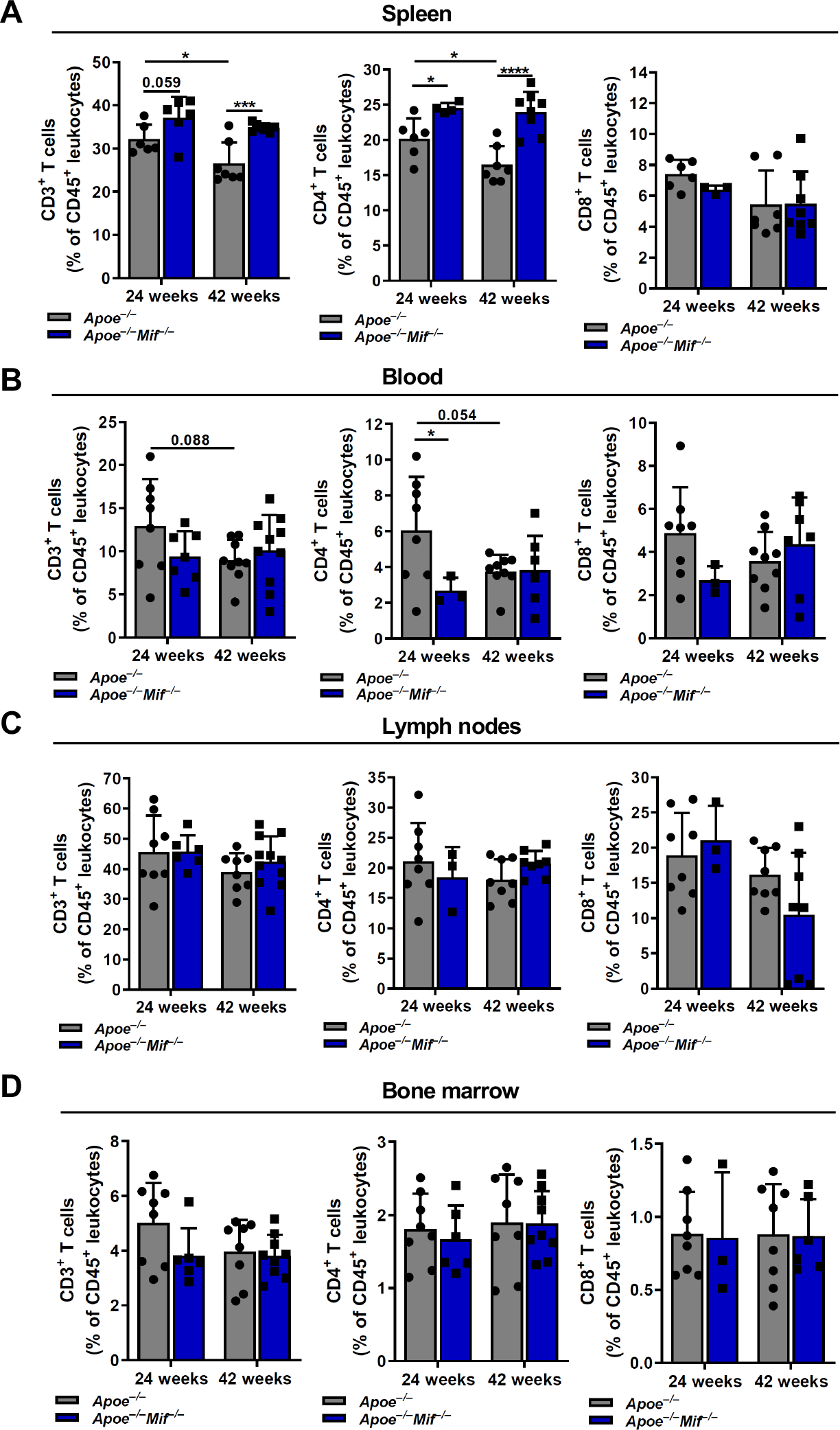
CD4^+^ T cell numbers in spleen and blood of hyperlipidemic *Apoe^−/–^* mice change in a MIF and age-dependent manner. FACS-based quantification of total CD3^+^ T cells (*left panels*), CD4^+^ T-cell subsets (*middle panels*), and CD8^+^ T-cell subsets (*right panels*) in spleen (**A**), blood (**B**), lymph nodes (**C**), and bone marrow (**D**) of *Apoe^−/–^Mif^−/–^* mice (blue) and comparison to *Apoe^−/–^* controls (grey). 30- and 48-week-old mice on HFD for 24 and 42 weeks, respectively, were examined. Bars shown represent data from n=3-8 independent mice and results are shown as means ± SD. Statistics: two-way ANOVA; *, *P*<0.05; ***, *P*<0.001; ****, *P*<0.0001, non-significant results with *P* values between 0.05 and 0.1 are indicated.

The T-cell analysis also revealed a significant influence of MIF. Splenic CD4^+^ T-cell numbers in *Apoe*^−*/*−^*Mif*^−/–^ mice were significantly increased compared to T-cell numbers in spleens from *Apoe*^−*/*−^ mice, both at 24 and 42 weeks of HFD, with a more pronounced difference seen at 42 weeks (**Figure 2a**). This effect in the aged 48/42-week mice was further confirmed in the 52/6-week aging/HFD model (**Supplementary Figure S1i**). To further explore whether the observed increase in splenic T-cell numbers in *Apoe*^−*/*−^*Mif*^−/–^ mice could contribute to the atheroprotection phenotype in these mice, we quantified FoxP3 mRNA levels as a marker for regulatory T cells (Tregs), which are considered anti-atherogenic. Splenic FoxP3 levels were elevated in *Apoe*^−*/*−^*Mif*^−/–^ mice after 42 weeks of HFD but not in the 24-week-HFD group (**Supplementary Figure S3**), suggesting that other mechanisms than deregulated Treg numbers underlie the lost atheroprotection in *Mif*-deficient mice upon aging. While no MIF-dependent changes were observed for T cells from LN or BM **(Figure 2c-d)**, blood CD4^+^ T-cell numbers were lower in *Apoe*^−*/*−^*Mif*^−/–^ mice compared to *Mif*-expressing controls in the 30/24- as well as in the 52/6-week group (**Figure 2b**, **Supplementary Figure S1j**).

We also determined B-cell, monocyte and neutrophil counts in spleen, LN, BM and blood. While monocyte counts were not altered between age groups or as a function of MIF, some changes were seen for B-cells and neutrophils such as a decrease of LN B cells in 24- week HFD-fed *Apoe*^−*/*−^*Mif*^−/–^ mice, an increase of CD19^+^ B cells in the blood and in the LNs of *Apoe*^−*/*−^*Mif*^−/–^ mice at 42 weeks HFD, or an increase in BM neutrophils in 24-week HFD-fed *Apoe*^−*/*−^*Mif*^−/–^ mice (**Supplementary Figure S4**). However, these changes were subtle and did not give rise to an apparent consistent pattern that could explain the observed plaque phenotype, and thus were not pursued further.

Together, the immune cell analysis in blood and peripheral lymphoid organs indicated that especially CD4^+^ T-cell numbers in the spleen may be controlled in a MIF- and age-dependent manner, but the determined increase in FoxP3^+^ Treg cells in the spleens of *Mif*- deficient mice in the 48/42- but not 30/24-week group does not explain the age-related loss of protection phenotype.

### Lesional Trem2^+^ macrophages decrease in global *Mif*-deficient *Apoe*^−*/*−^ mice over the course of aging

To further investigate the loss of atheroprotection in aged *Mif*-deficient mice, we analyzed SMCs and macrophages in plaques of the BCA after 24 and 42 weeks of HFD. While the content of smooth muscle actin (SMA)^+^ SMCs was unaltered (**Figure 3a-b**), we detected substantial changes in plaque macrophage numbers (**Figure 3c-d**). As indicated by vessel staining with an anti-CD68 antibody, macrophage numbers in plaques of *Apoe*^−*/*−^*Mif*^−/–^ mice fed an HFD for 24 weeks were almost doubled compared with those in *Apoe*^−*/*−^ mice; intriguingly, this increase was gone in the 42-week HFD regimen. This was mainly due to a significant decrease in macrophages numbers in the *Apoe*^−*/*−^*Mif*^−/–^ mice over the course of aging (**Figure 3c-d**). As this effect paralleled the observed loss of protection from plaque formation in the *Apoe*^−*/*−^*Mif*^−/–^ mice between 24 and 42 weeks of HFD, we hypothesized that the stage-dependent decrease in macrophages could be predominantly due to a reduction in an atheroprotective macrophage subset. To this end, single cell (sc) RNAseq analysis recently revealed an accumulation of lipid-loaded ‘foamy’ Trem2^hi^ macrophages within atherosclerotic lesions, a previously unrecognized macrophage subtype assigned anti-inflammatory and homeostatic functions (35–37). We therefore next analyzed our plaque specimens for Trem2^+^ macrophages and found significantly reduced Trem2 expression in plaque areas of *Mif*- deficient *Apoe*^−*/*−^ mice after 42 weeks of HFD compared to the 24-week HFD group (**Figure 3e-f; Supplementary Figure S5**). The additional determination of Trem2^+^ Ki67^+^ cells showed that these proliferating Trem2^+^ cells were increased in the 24-week *Mif*-deficient *Apoe*^−*/*−^ mice compared to wildtype control, while Trem2^+^ Ki67^+^ cells were decreased in the *Mif*-deficient *Apoe*^−*/*−^ mice after 42 weeks of HFD compared to the 24-week group (**Figure 3g-h; Supplementary Figure S5**). This indicated that *Mif* deficiency may have a pro-proliferative effect on Trem2^+^ cells that is lost in mice of a more advanced age.

**Figure 3:**
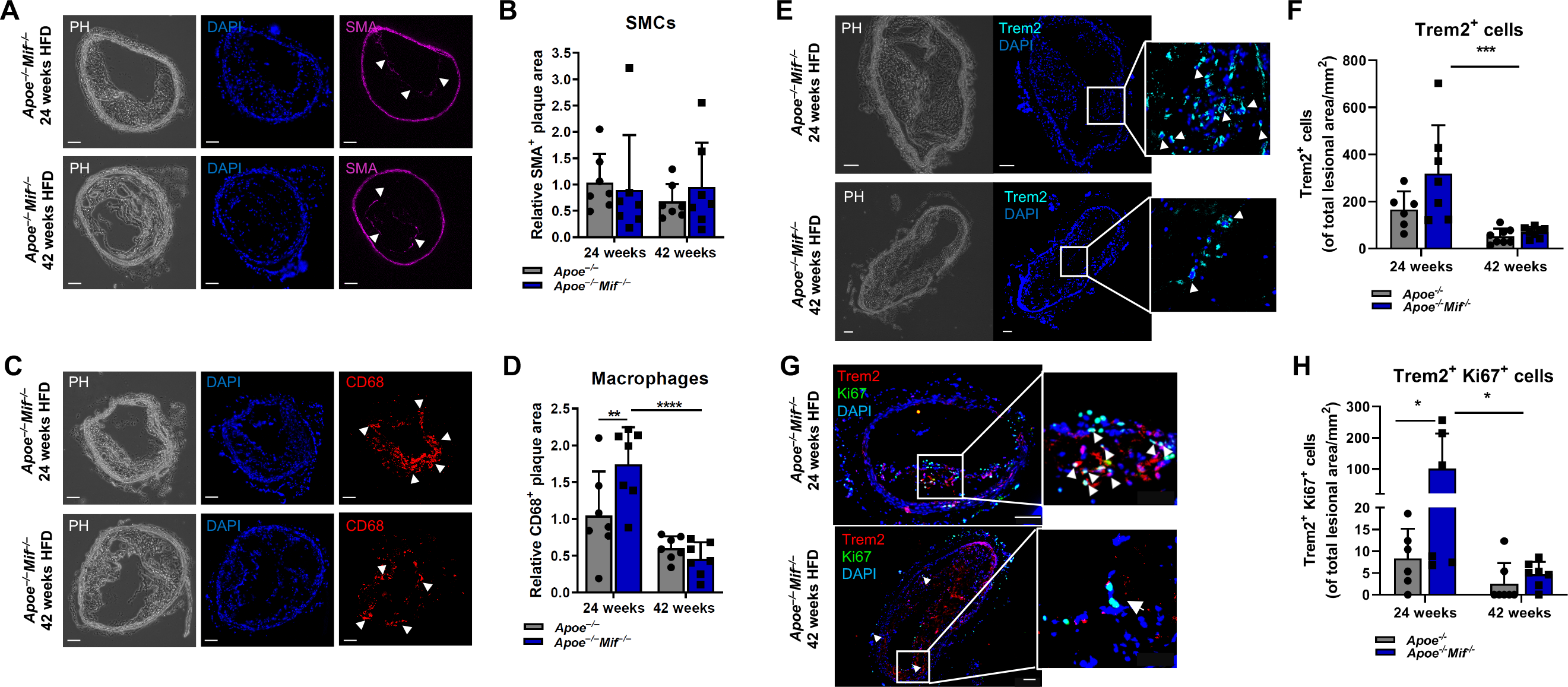
Age- and MIF-dependent changes in lesional total macrophage and Trem2^+^ macrophage numbers. Immunohistochemistry was performed on BCA sections derived from 30- and 48-week-old *Apoe^−/–^Mif^−/–^* (blue) and *Apoe^−/–^* (grey) mice after 24 and 42 weeks of HFD, respectively. Smooth muscle cells (SMA), total macrophage content (CD68), and Trem2^+^ macrophages were quantified in the plaque area. Nuclear counterstain was performed using DAPI. **A-B**) Representative images (**A**) and quantification (**B**) of the SMA^+^ plaque area. 2-3 sections were analyzed per mouse and mean values normalized to values of the 30-week-old *Apoe^−/–^* controls that received HFD for 24 weeks (n=7 mice; results presented as means ± SD; scale bar: 100 µm; arrows indicate the SMA^+^ plaque area; PH=phase contrast). **C-D**) Representative images (**C**) and quantification (**D**) of the CD68^+^ plaque area. Analysis performed as in (**A-B**) (n=7, results are presented as means ± SD; scale bar: 100 µm; arrows indicate the CD68^+^ plaque area). **E-F**) Representative images (**E**) and quantification (**F**) of Trem2^+^ cells within the plaque. Cells are indicated by arrow heads in the enlarged image sections. Trem2^+^ cells were quantified manually and are depicted as Trem2^+^ cells per mm^2^ plaque area. As Trem2 is mainly expressed by myeloid cells, the stained cells were considered as Trem2^+^ macrophages. **G-H)** Representative images (**G**) and quantification (**H**) of lesional Trem2^+^ Ki67^+^ cells. Trem2^+^ Ki67^+^ cells are indicated by arrow heads in the enlarged image sections. Trem2^+^ Ki67^+^ double-positive cells were manually quantified and are depicted as Trem2^+^ Ki67^+^ cells per mm^2^ plaque area. Results are presented as means ± SD from n=6-8 mice; scale bar: 100 µm. Statistics: two-way ANOVA; *, *P*<0.05; ***, *P*<0.001; each data point represents one independent mouse.

Together, the analysis of lesional immune cells in the BCA suggested that age/HFD duration-dependent loss of atheroprotection in *Apoe*^−*/*−^*Mif*^−/–^ mice is associated with a reduction in anti-inflammatory Trem2^+^ plaque macrophages.

### *Mif*-deficiency accelerates formation of stage I/II ATLO-like lymphocyte-rich cell clusters while ATLO numbers match in *Mif*-deficient and wildtype mice at advanced aging stage

While we concluded that the above-described change in plaque Trem2^+^ macrophages contributes to the age-dependent attenuation-of-atheroprotection phenotype in *Apoe*^−*/*−^*Mif*^−/–^ mice, we surmised that additional cellular MIF-driven mechanisms may be involved as well. B cells are rare in atherosclerotic lesions but are present in the adventitia and adjacent periadventitial tissue (38). Since we recently uncovered a link between MIF and B cells in atherosclerotic lesions (23), we next determined peri-adventitial cell clusters in the BCA over the course of aging.

After 24 weeks of HFD, we observed peri-adventitial cellular clusters, reminiscent of lymphocyte/B cell-rich clusters, in the BCA of *Apoe*^−*/*−^*Mif*^−/–^ mice, while such cell clusters were rare in age-matched *Apoe*^−*/*−^ controls (**Figure 4a-b**). Interestingly, this difference in cluster formation disappeared in aged mice that received HFD for 42 weeks, as the number of clusters increased in *Apoe*^−*/*−^ mice at this aging stage (**Figure 4b**). Thus, *Mif*-deficiency accelerates cluster formation in *Apoe*^−*/*−^ mice in medium-aged hyperlipidemic mice, but this difference is lost towards more advanced aging stages. We hypothesized that the clusters contribute to the atheroprotective effect of *Mif*-deficiency after 24 weeks of HFD and therefore characterized their nature and cellular composition. Adventitial immune cell aggregates have been predominantly discussed in the context of sustained chronic inflammation and the formation of ATLOs in aged atherogenic *Apoe^−/–^* mice (28). Thus, we determined whether the identified cellular clusters in *Mif*-deficient mice could represent peri-adventitial ATLOs. Due to their T- and B-cell-rich lymphoid organ-like structure, ATLOs can be identified through a variety of markers, including those for lymphocytes, lymph vessels, high-endothelial venules (HEVs), and capsule components. Using anti-CD3e and anti-B220 antibodies, we detected the presence of T and B cells, respectively, in peri-adventitial clusters of *Apoe*^−*/*−^*Mif*^−/–^ mice (**Figure 4c**). Moreover, the clusters contained lymph vessels and conduit-like structures, as evidenced by pronounced staining for Lyve-1 and ER-TR7, respectively, and contained substantial collagen type IV positivity (**Figure 4c**), together suggesting that the clusters display an ATLO-like composition. However, features of mature ATLOs were absent, as indicated by lack of staining for peripheral node addressin (PNAd), a marker of HEVs, and CD35, a marker for follicular dendritic cells (FDCs) (**Figure 4c**). Together, these results suggested that the prominent lymphocyte-rich clusters that we detected in the BCA of 24-week-HFD-fed *Apoe*^−*/*−^ *Mif*^−/–^ mice as well as in the BCA of aged 42-week-HFD-fed *Apoe*^−*/*−^*Mif*^−/–^ and *Apoe*^−*/*−^ mice likely represent *early* ATLOs, corresponding to a stage I or II ATLO (29).

**Figure 4:**
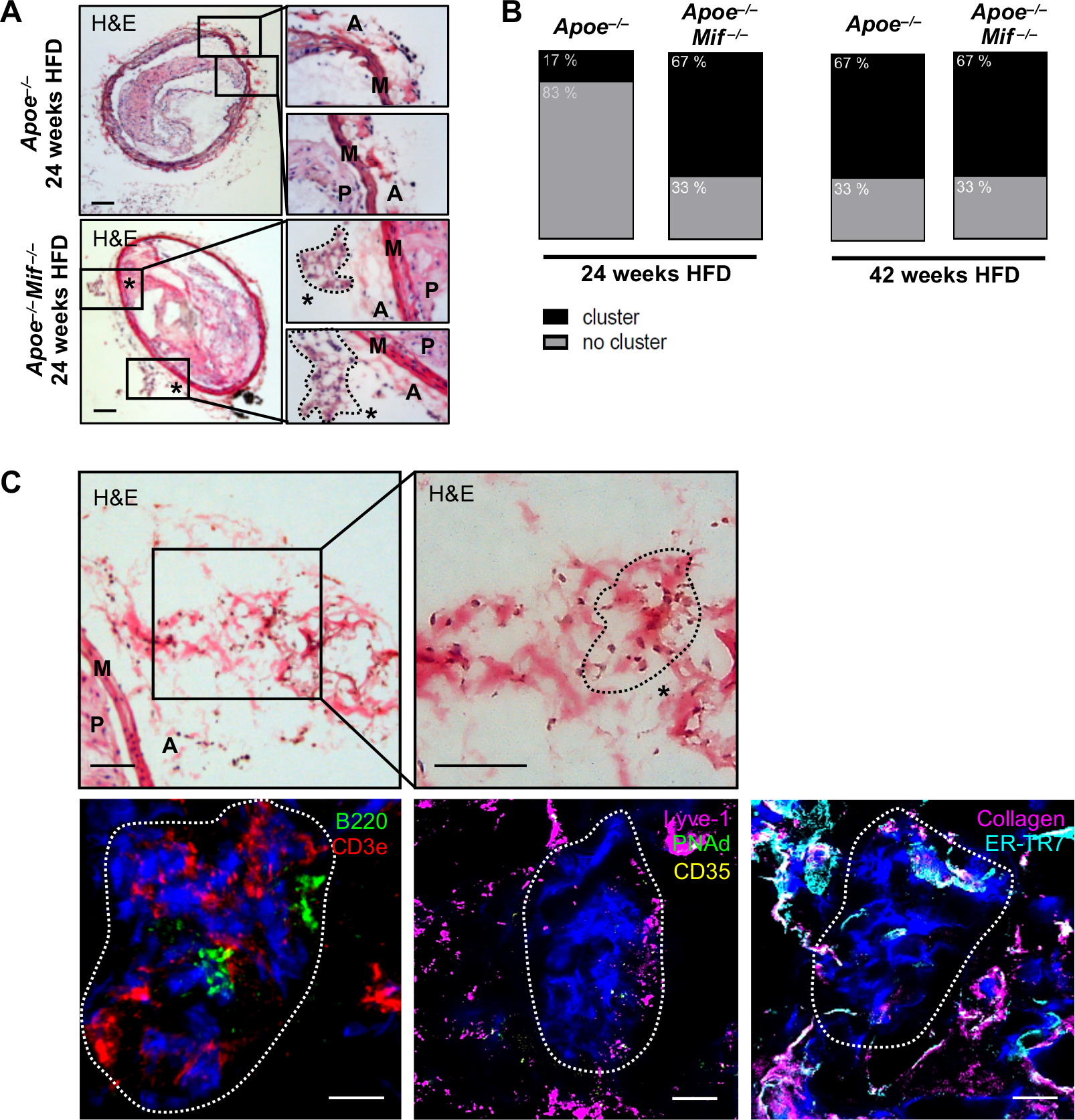
*Mif*-deficiency accelerates the formation of lymphocyte-rich stage-I/II (*early*) ATLOs. **A-B)** Detection of lymphocyte-rich (peri-)adventitial cell clusters in the BCA of *Mif*- deficient versus *Mif*-expressing hyperlipidemic *Apoe^−/–^*mice. **A)** Representative images of lymphocyte-rich (peri-)adventitial cell clusters in the BCA of 30-week-old *Apoe^−/–^* control mice (upper panel) *versus Apoe^−/–^Mif^−/–^* mice (lower panel) on HFD for 24 weeks as assessed by H&E-staining. A total of 10 serial sections with a distance of 50 µm was screened for lymphocyte-rich (peri-)adventitial cell clusters for each mouse. Boxed areas containing lymphocyte-rich (peri-)adventitial cell clusters (lower panel) or no clusters (upper panel) are also shown at higher magnification (right hand panels). Cell clusters in the boxed areas of the lower panel are indicated by an asterisk and circled in the enlarged panels. A=adventitia, M=media, P=plaque, scale bar: 100 µm. **B**) Quantification of the detected lymphocyte-rich (peri-)adventitial cell clusters comparing *Apoe^−/–^Mif^−/–^* and *Apoe^−/–^* mice at both the 30/24 and 48/42 age/HFD time intervals (n=6 mice per group). **C**) The lymphocyte-rich (peri-)adventitial cell clusters in *Mif*-deficient *Apoe^−/–^* mice are stage I/II ATLOs. Representative images of the immunostaining of the clusters against ATLO-specific markers (lower panel) including B220 (B cells), CD3e (T cells), LYVE-1 (lymphatic vessel endothelial hyaluronic acid receptor 1; lymph vessels), PNAd (peripheral node addressin; high endothelial venules), CD35 (follicular dendritic cells), ER-TR7 (reticular fibers/fibroblasts), and collagen IV (scale bar: 10 µm). For a better visualization of the localization of the clusters, the H&E images of the respective region are shown (upper panel). The circled region in the enlarged H&E image section marked by an asterisk indicates the analyzed cluster (scale bar: 50 µm).

### A decrease in anti-oxLDL IgM antibody titer correlates with attenuated atheroprotection at more advanced aging stages

While the pattern of ATLO formation across aging stages can only partially explain the loss of atheroprotection observed in aged *Mif*-deficient *Apoe*^−*/*−^ mice, the increase in numbers of early ATLOs in the BCA of 24-week-HFD-fed *Apoe*^−*/*−^*Mif*^−/–^ mice could well contribute to atheroprotection seen in *Mif*-deficient mice at this age, as ATLO-derived B cells have been attributed atherogenesis-dampening properties (30, 39).

To study this possibility, we analyzed the plasma levels of atherosclerosis-relevant anti-oxLDL antibody isotypes (40). Anti-oxLDL IgM antibody levels that have been associated with atheroprotection (40) were significantly decreased after 42 weeks of HFD in *Apoe*^−*/*−^*Mif*^−/–^ mice (**Figure 5a**). In line with these results, titers of the anti-oxLDL IgG isotype, known to promote atherosclerosis (40), were not found to be decreased (**Figure 5b**). We conclude that the observed age-dependent decrease of anti-oxLDL antibodies of the IgM isotype could contribute to the attenuation of atheroprotection seen in 42-week-HFD-fed *Apoe*^−*/*−^*Mif*^−/–^ mice.

**Figure 5:**
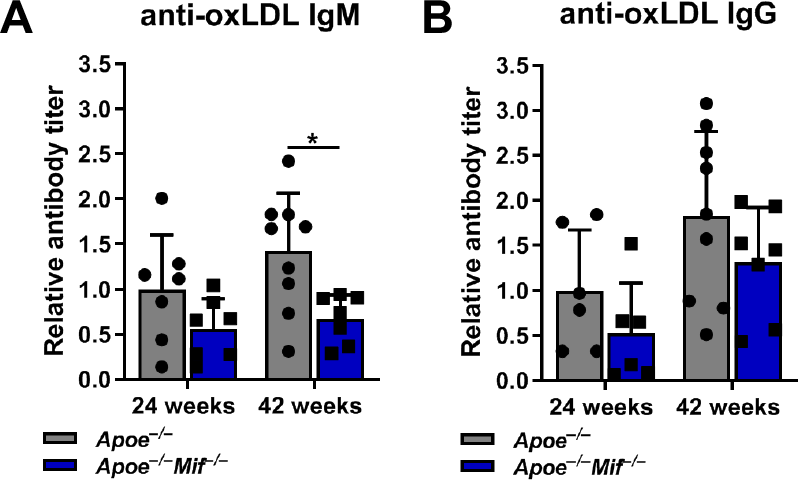
*Mif*-deficiency leads to a decrease in atheroprotective anti-oxLDL IgM antibody titers in highly-aged hyperlipidemic *Apoe^−/–^*. Plasma levels of anti-oxLDL IgM (**A**) and anti- oxLDL-IgG (**B**) antibodies in 30- and 48-week-old *Apoe^−/–^Mif^−/–^* mice (blue) *versus Apoe^−/–^* controls (grey) on 24- and 42-week-HFD, respectively. oxLDL-coated microtiter plates were incubated with mouse plasma and HRP-labelled anti-IgM/IgG antibodies and chemi- luminescent signals obtained. Values are normalized to those of the 30/24-week *Apoe^−/–^* mouse group. Data are from n=6-9 mice and results are presented as means ± SD. Statistics: two-way ANOVA; *, *P*<0.05; each data point represents one mouse.

## Discussion

Macrophage migration inhibitory factor (MIF) is an inflammatory cytokine and atypical chemokine that was previously shown to promote atherogenesis by enhancing CXCR2/4- mediated leukocyte recruitment and vascular inflammation. The current study for the first time establishes a link between MIF and age-related changes in atherosclerotic pathology, demonstrating that regio-specific atheroprotection in global *Mif*-gene-deficient atherogenic *Apoe*^−*/*−^ mice is lost during the course of aging and Western-type HFD. As potential underlying cause, we identify a combination of mechanisms, most notably an age-dependent reduction in lesional Trem2^+^ macrophages in the absence of MIF, but also a loss in relative abundance of lymphocyte-rich-stage I/II ATLOs, and decreased levels of circulating anti-oxLDL IgM antibodies in *Mif*-deleted aged atherogenic *Apoe*^−*/*−^ mice compared to *Mif*-proficient controls. **Figure 6** summarizes these mechanisms and their stage-dependent effects on athero- protection and *Mif*-deficiency.

**Figure 6:**
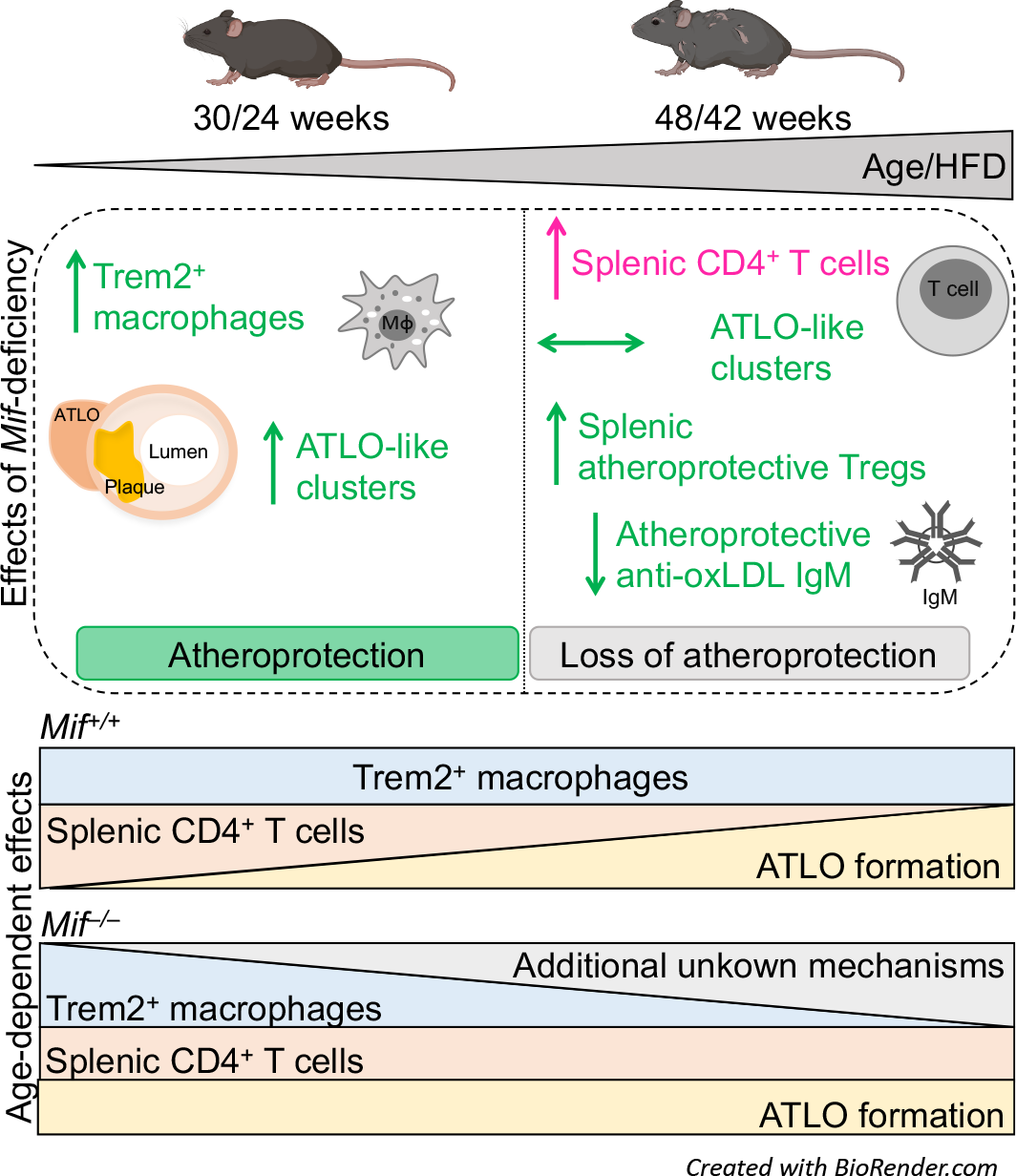
Scheme summarizing the age- and MIF-dependent effects in advanced atherosclerosis. The scheme summarizes the effects of global *Mif* gene-deficiency in 30- and 48-week-old *Apoe^−/–^* mice that received HFD for 24 and 42 weeks, respectively. The atheroprotective phenotype of *Mif* gene deletion in the 30/24-week group was accompanied by an increase in lesional Trem2^+^ macrophages and accelerated atheroprotective ATLO-like cluster formation in comparison to *Mif*-proficient *Apoe^−/–^* controls. Atheroprotection was lost in the 48/42-week group, accompanied by lowered atheroprotective anti-oxLDL IgM antibody levels and increased splenic CD4^+^ cells in the *Mif*-deficient animals, while atheroprotective Tregs were increased in the spleen of these mice. Additionally, no differences in atheroprotective ATLO-like cluster formation between *Mif^−/–^Apoe^−/–^* and *Apoe^−/–^* mice were observed in the 48/42-week group. These effects resulted in an overall loss of atheroprotection in the aged mice. Atheroprotective mechanisms are depicted in green, while atheroprogressive ones are shown in magenta. HFD, high-fat diet; Trem2, triggering receptor expressed on myeloid cells 2; ATLO, artery tertiary lymphoid organ; oxLDL, oxidized low density lipoprotein. The mouse icons were generated with BioRender.com.

MIF has been attributed an overall pro-inflammatory and atheroprogressive role as evidenced in various experimental models of atherosclerosis applying *Mif*-gene deficiency or antibody blockade and in line with correlations between MIF expression levels and human atherosclerotic disease (8, 10, 18, 20, 22). However, while the specific role of the cardiac MIF/AMPK axis was examined in ischemic recovery in the senescent heart (34), the contribution of MIF to age-driven exacerbation of atherosclerotic lesion formation has not been systematically studied. Here we analyzed the influence of *Mif*-gene deficiency in *Apoe*^−*/*−^ mice across different ages and HFD exposure intervals. *Mif*-knockout and wildtype *Apoe*^−*/*−^ mice were compared at an age of 30, 42, and 48 weeks, corresponding to 24, 36, and 42 weeks of HFD treatment, respectively. To further study the influence of aging *versus* duration of exposure to a cholesterol-rich HFD, a model of 52-week-old mice, in which HFD was limited to the last 6 weeks, was employed.

Deletion of the *Mif*-gene in the atherogenic *Ldlr*^−*/*−^ mouse model leads to protection from atherosclerotic plaque formation across the entire vascular bed, including aortic root and arch (18, 21). In contrast, protection conveyed by global *Mif*-deletion in the context of the *Apoe*^−*/*−^ model is regio-specific and limited to the BCA and abdominal aorta (23). The precise mechanistic causes for this background-specific difference have yet to be explored, and it is currently unclear whether a similar phenotype would be seen in conditional, cell/tissue-specific models of *Mif*-gene deletion, such as in arterial- or myeloid-specific *Mif*-knockouts. Here, we confirm the regio-specific atheroprotective effect of *Mif*-deficiency in BCA and abdominal aorta in 30-week-old *Apoe*^−*/*−^ mice on HFD for 24 weeks. Notably, while 48-week-old *Mif*-deficient mice on HFD for 42 weeks showed significantly reduced serum triglyceride levels compared to age-matched *Mif*-proficient control animals, the site-specific atheroprotective effect was reduced or “lost” in these mice, as well as in 52-week-old mice that had been on HFD for the last 6 weeks. As this ruled out major metabolic effects as an underlying cause for the observed effect, we surmised that *Mif*-deficiency unleashes suppressed atheroprotective mechanisms (Trem2^+^ macrophages, ATLOs), properties of MIF that are lost upon aging. The latter could be due to an overall increased inflammatory status in the aged atherosclerotic mice. In fact, inflammatory cytokine levels such as those of interleukin-6 (IL-6) or IL-1β have been found to increase over the course of aging and have been suggested to contribute to aging-related exacerbation of atheroprogression as both vascular extrinsic and intrinsic factors (41–43). Regarding MIF, the correlation between the regulation of MIF expression and aging has remained unclear, but one study in humans suggests an increase in circulating MIF levels in the elderly (44), whereas age-related tissue expression of murine MIF was found to be increased in aorta (45) but decreased in cardiac tissue (34). We determined MIF mRNA expression levels in spleen and liver and MIF protein levels in plasma in our mouse model, but did not detect any differences between the 24- and 42-week-HFD groups. Thus, age-related changes in MIF expression cannot readily explain the observed loss of atheroprotection in the aged *Mif*-deficient animals.

We noted a clear-cut regulatory effect of MIF on macrophage accumulation in BCA lesions, which was lost upon aging. Compared to *Apoe*^−*/*−^ controls, *Mif*-deficient *Apoe*^−*/*–^ mice exhibited higher lesional macrophage counts in 30-week-old animals after 24 weeks of HFD, but no difference was apparent in the aged 48/42-week mice. This was mainly due to a marked drop in lesional macrophage numbers in the *Mif*-deficient mice over the course of aging. Interestingly, sub-class-specific analysis indicated that the number of Trem2^+^ macrophages, a subtype deemed anti-inflammatory, significantly decreased between the 30/24 and 48/42 time points, suggesting that it could, at least partially, account for the age-dependent loss of atheroprotection in *Mif*-gene-deficient mice. Of note, arterial immune cell infiltration/content as a vascular intrinsic factor has been linked to the effects of aging on vascular inflammation and lesion formation (41, 46, 47). Using an *Ldlr^−/–^* model of atherosclerosis, it was additionally shown that aging in combination with HFD induces monocytosis and aortic macrophage accumulation (48). How aging affects immune cell numbers specifically in the BCA of *Apoe*^−*/*−^ mice has not been investigated. Our data suggest that arterial MIF could serve as a regulator of immune cell homeostasis in the BCA, suppressing – either directly or indirectly - homeostatic macrophage sub-populations such as the Trem2^+^ subtype. Our quantification of Trem2^+^ Ki67^+^ cells indicates that this could be due to an effect on the proliferative capacity of lesional Trem2 ^+^ cells. Of note, Trem2^+^ macrophages were recently associated with anti-inflammatory properties and plaque regression (36, 37). Accordingly, MIF in the vasculature might contribute to an imbalance between macrophage subtypes, overall promoting atherosclerotic plaque formation, a property that is lost or overridden in the course of aging. MIF may thus display a dichotomic and biphasic role in the atherogenic vasculature. While it primarily promotes leukocyte infiltration and vascular inflammation in young and middle-aged mice through its chemokine-like inflammatory activities as shown in numerous studies (10, 18, 20, 21, 33), and may attenuate the proliferation or survival of anti-inflammatory plaque macrophage subtypes; these effects may be lost or become overcompensated in the aged vessel wall. Alternatively, MIF may have a polarizing effect on plaque macrophages from an anti- to a pro-inflammatory phenotype, as observed previously in the context of oncogenesis or adipose tissue-associated macrophages (49, 50). A detailed future investigation of MIF effects on the recruitment, survival, or polarization of selective vascular wall macrophage sub-populations, including the involved MIF receptor pathways, would thus be warranted.

Interestingly, we also observed an influence of MIF on T cells. Splenic CD4^+^ T-cell numbers in *Apoe*^−*/*−^*Mif*^−/–^ mice were significantly increased both at 24 and 42 weeks of HFD compared to T-cell numbers in spleens from *Apoe*^−*/*−^ mice. Due to the decrease of T-cell numbers in *Apoe*^−*/*−^ mice between 24 and 42 weeks of HFD, this difference was particularly pronounced at 42 weeks HFD. It is well established that aging affects the anatomy and organization of the spleen (51). Different studies revealed a lowered abundance of splenic T-cell zones in aged C57BL/6 mice that was accompanied by a reduction in CD4^+^ T-cell recruitment and function (16, 52). We noticed that peripheral CD4^+^ T-cell counts were significantly decreased in *Mif*-deficient mice after 24 weeks of HFD, in line with our earlier findings (23), as well as in the 52/6-week model of aging and hyperlipidemia. This could indicate that *Mif*-deficiency might promote T-cell egress from the periphery and favor homing to the spleen. However, the mechanism for this scenario remains unclear and our analysis of splenic Treg numbers did not reveal an obvious connection between splenic T-cell subtypes and the observed plaque phenotype in the *Mif*-deficient animals.

We previously observed an enhanced re-organization of B cells into cluster-like structures in peri-adventitial areas in proximity to atherosclerotic plaques in the BCA of *Mif*- deficient *Apoe*^−*/*−^ mice (23). Here, we show that the formation of lymphocyte-rich clusters in *Apoe*^−*/*−^ mice is age-dependent and that *Mif*-deficiency accelerated the formation of the clusters at 24 weeks of HFD, an effect that paralleled the attenuation of lesion formation due to *Mif*- deficiency. We also noted that the increase in cluster formation in *Mif*-deficient mice compared to *Mif*-expressing controls vanished during the course of aging. The nature of these clusters had been elusive. Here, we provide evidence that the clusters that form in an accelerated manner in *Mif*-deficient *Apoe*^−*/*−^ mice are stage I/II or early stage ATLOs. Adventitial ATLOs are typically observed in highly aged *Apoe*^−*/*−^ mice on chow or HFD for 52-78 weeks. ATLOs display a lymphoid-like organ structure and ATLO-based lymphocytes are found in complex immune structures with organized T- and B-cell zones including germinal centers (GCs), lymph vessels, high-endothelial venules, and follicular dendritic cells (FDCs) (29, 53). Based on the expression of various markers, ATLOs of different maturation stages, i.e. stage I, II, and III, can be classified (29, 54). Moreover, ATLOs need to be differentiated from fat-associated lymphoid clusters (FALCs), which are T/B-cell aggregates that are exclusively observed in adipose tissue (29). Stage I ATLOs feature loosely arranged B and T cells and lymphocytes in these early ATLOs are not organized in dedicated B- and T-cell areas. They are also characterized by the expression of lymphorganogenic chemokines such as CXCL13. Stage II ATLOs contain separate T- and B-cell areas and lymph vessel neogenesis becomes prominent together with a dense network of lymph node-like conduits connecting the arterial wall with newly formed HEVs. In contrast, fully matured or stage III ATLOs display separate T-cell and B-cell follicles, activated germinal centers (GCs) with FDCs, plasma cell (PC) niches, lymph vessels, as well as blood vessel neogenesis. In addition to an accumulation of B and T cells, we detected a vascular system including LYVE-1^+^ lymphatic vessels, and appreciable collagen IV expression. However, organization of the infiltrated lymphocytes into clearly distinct T- and B-cell zones as well the formation of GCs with FDCs that is characteristic of stage III ATLOs were absent (29). Overall, the features are indicative of stage I or II ATLOs and we hypothesize that they may develop into mature stage III ATLOs upon further aging. As our structures only contained sparse ER-TR7 positivity, we conclude that they are distinct from secondary lymphoid organs (SLO), which would exhibit a well-structured ER-TR7+ encapsulation (29).

The functional role of ATLOs in atherosclerosis is incompletely understood. However, recent data revealed an atheroprotective role accompanied by activation of anti-inflammatory Treg cells within the ATLO (39). Moreover, different B-cell subsets have been described to be present in ATLOs with B1b B cells representing the majority of the B-cell compartment (30). B1 cells have been attributed atheroprotective functions by secreting natural IgM (55, 56). Based on these findings, we hypothesize that our identified stage I/II ATLOs may contribute to the atheroprotective effect in *Mif*-deficient *Apoe*^−*/*−^ mice that is eventually attenuated during aging. Accordingly, we next sought to determine if the formation of stage I/II ATLOs was accompanied by changes in the production of atheroprotective IgM *versus* atheroprogressive IgG antibodies. The mechanism by which IgG and IgM promote or attenuate atherosclerosis are mainly based on the recognition of oxidation-specific epitopes (OSEs) such as in oxLDL (40). Atheroprotective anti-oxLDL IgM antibodies have been shown to neutralize oxLDL and inhibit its uptake by macrophages thereby suppressing foam cell formation, whereas anti-oxLDL IgG antibodies form complexes with oxLDL that promote lesional macrophage activation via Fcγ-receptors (40). Here, we noted that loss of atheroprotection in the more- aged *Mif*-deficient mice correlated with reduced levels of circulating atheroprotective anti-oxLDL IgM levels, whereas anti-oxLDL IgG levels remained unchanged.

Our study of *Mif*-deficiency in the context of atherosclerosis and aging furthers the notion that aging affects atherogenesis in a complex manner, with cytokine/chemokine-related inflammatory mechanisms playing key roles. Considering the continuing rise of atherosclerotic patients in an aging population, the design of treatment strategies specifically tailored for elderly individuals will become important. Based on the principal success of the CANTOS trial (57) and the emerging link between inflamm’aging and atherosclerosis (25), cytokine-/chemokine-directed strategies may represent promising approaches for age-tailored intervention strategies. MIF is a promising target in atherosclerosis and a variety of inhibition routes including antibody-, small molecule- and peptide-based strategies have been discussed (9, 58, 59). However, age-tailored strategies have not been investigated. The preclinical data obtained in our present study confirm numerous previous studies suggesting that MIF-based inhibition strategies may be promising in cardiovascular patients. The ‘attenuated- atheroprotection phenotype’ related to *Mif*-deficiency and advanced aging was observed in global *Mif*-gene-deficient mice. Global gene deficiency may cause various compensatory mechanisms and MIF has been assigned both extracellular inflammatory and intracellular homeostatic activities (7). Moreover, the MIF-homolog D-dopachrome tautomerase (D- DT)/MIF-2 shares some activities with MIF (60), while distinct effects have also been noted (50). MIF-2 has not been studied in atherosclerosis, but (compensatory) effects of MIF on MIF- 2 gene expression have been observed in a cancer cell line (61). It would thus be interesting to test whether the observed ‘attenuated-atheroprotection phenotype’ seen in the context of global *Mif*-deficiency, is recapitulated in conditional cell/tissue-specific *Mif*-gene-deficient models, or upon MIF depletion by a pharmacological strategy. Furthermore, it may be argued that general MIF-blocking strategies might be sub-optimal in highly-aged individuals and that instead, receptor pathway-specific MIF-targeting strategies should be considered (58).

Such conclusions may be hampered by the well-known challenges of translating data from mouse models into human disease scenarios. To this end, potential specific limitations of the current study are: i) the aforementioned use of global *Mif*-gene-deficient mice; ii) the confinement to the *Apoe*^−*/*−^ mouse model with its regio-specific phenotype that has not been observed in *Mif*^−*/*−^*Ldr* ^−*/*−^ mice (18, 21); iii) the according comparison between BCA and aortic root or arch lesions; iv) and partly limited animal numbers and no confirmation in a second independent set of cohorts, owing to the long study times and 3R rule considerations. On the other hand, clinical correlation studies in CVD patients taking into account MIF expression levels in *MIF*-low (carrying the CATT5/5 or 5/6 polymorphism) *versus MIF*-high (carrying the CATT6/7 or 7/7 polymorphism) expressers (62, 63) could elegantly guide translational considerations.

In conclusion, this work has unravelled a previously unknown link between *Mif*- deficiency, atheroprotection, and aging in the *Apoe*^−*/*−^ model of atherosclerosis. We show that site-specific atheroprotection in global *Mif*-deficient mice is attenuated/lost in aged animals and/or upon long-term HFD and identify a combination of effects, encompassing reduced lesional Trem2^+^ macrophages, peri-adventitial ATLOs, and decreased atheroprotective anti-oxLDL IgM antibodies, as potential underlying causes of this phenotype. While the mechanistic details of these effects such as the involved receptor or interim pathways will have to be subject to future scrutiny, our study provides valuable molecular and cellular information that could assist in devising tailored strategies of MIF-directed targeting in atherosclerosis.

## Acknowledgements

This work was supported by Deutsche Forschungsgemeinschaft (DFG) grant SFB1123-A3 to J.B. and A.K., SFB1123-A1 to C.W., SFB1123-C2 to S.M., SFB-TRR219-M05 to H.N., and by DFG under Germany’s Excellence Strategy within the framework of the Munich Cluster for Systems Neurology (EXC 2145 SyNergy—ID 390857198) to J.B. and C.W., and LMUexc strategic partnerships to J.B. C.W. is Van de Laar Professor of Atherosclerosis. We thank Simona Gerra and Priscila Bourilhon for technical support; Christian Haass, Kai Schlepckow, Xianyuan Xiang, and Clement Cochain for providing us with Trem2 antibody; Marlies Zarwel for helpful discussions; and Sabrina Pagano and Sabine Steffens for advice regarding anti-oxLDL immunoassays. We are grateful to the mouse core facility of the Center for Stroke and Dementia Research (CSD) for their invaluable support with the mouse studies.

## Animal ethics approval

Animal experiments were approved by the local authorities (animal ethics approval ROB-55.2- 2532.Vet_02-18-040 of the government of Bavaria, Germany) and were performed according to the German animal protection law.

## Supplementary Material

### Supplementary Tables

**Supplementary Table S1:**
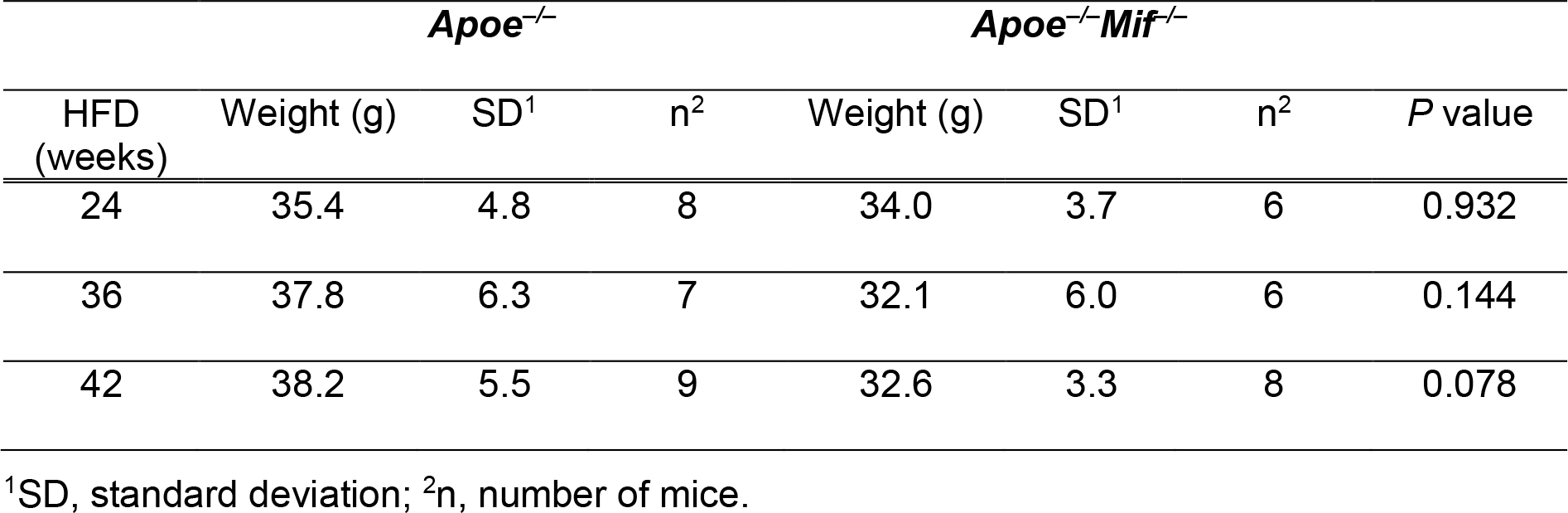
Body weights of the *Apoe–/–Mif–/–* and *Apoe–/–* mice after 24, 36, and 42 weeks of high-fat diet (HFD).

**Supplementary Table S2:** Plasma cholesterol and triglyceride levels of the *Apoe–/–Mif–/–* and *Apoe–/–* mice after 24 and 42 weeks of high-fat diet (HFD).

| HFD<br>(weeks) | Parameter | *Apoe*<sup>-/-</sup> |  |  | *Apoe*<sup>-/-</sup>*Mif*<sup>-/-</sup> |  |  | *P* value |
| --- | --- | --- | --- | --- | --- | --- | --- | --- |
|  |  | Mean | SD<sup>1</sup> | n<sup>2</sup> | Mean | SD<sup>1</sup> | n<sup>2</sup> |  |
| 24 | Cholesterol<br>(mg/dl) | 899.6 | 423.5 | 7 | 670.1 | 274.9 | 7 | 0.296 |
| 42 | Cholesterol<br>(mg/dl) | 1050.4 | 278.3 | 9 | 806.9 | 186.7 | 8 | 0.197 |
| 24 | Triglyceride<br>(mg/dl) | 132.8 | 68.6 | 7 | 134.4 | 59.5 | 6 | 0.999 |
| 42 | Triglyceride<br>(mg/dl) | 211.4 | 88.2 | 7 | 116.9 | 28.9 | 7 | 0.025 |

<sup>1</sup>SD, standard deviation; <sup>2</sup>n, number of mice.

**Supplementary Table S3:**
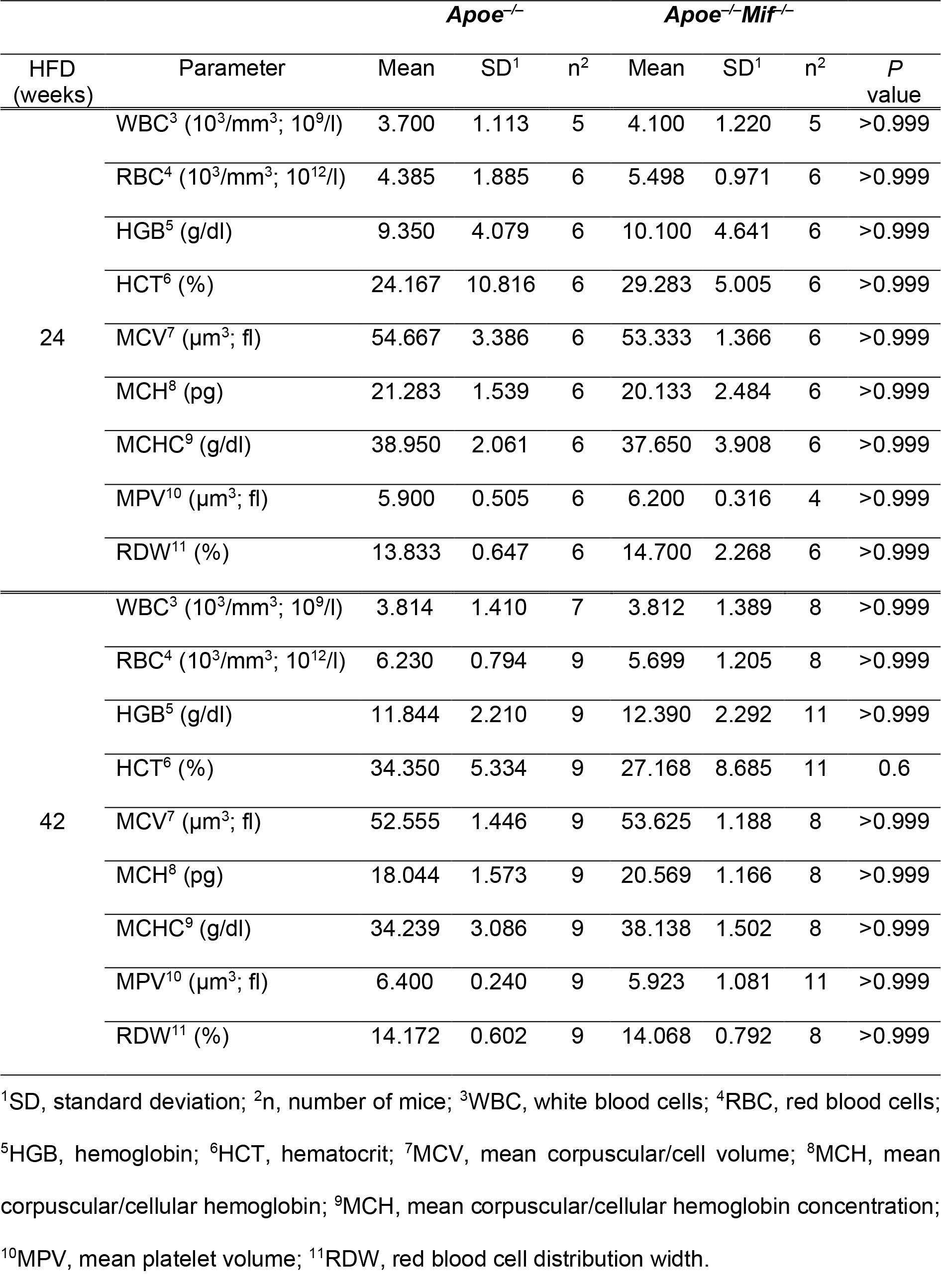
Blood parameters of *Apoe^−/–^Mif^−/–^* and *Apoe^−/–^* mice after 24 and 42 weeks of high-fat diet (HFD).

### Supplementary Figures

**Supplementary Figure S1:**
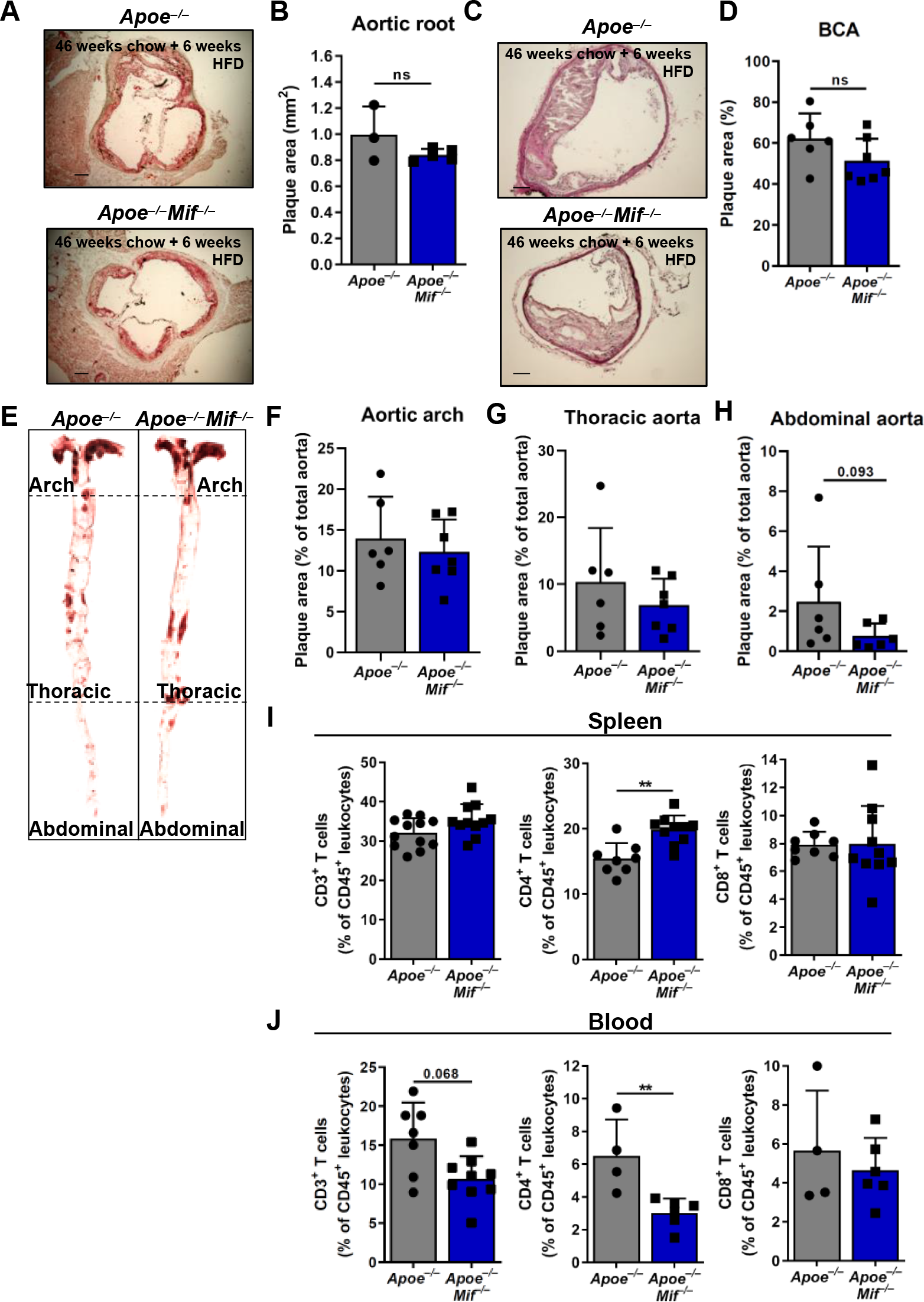
*Mif*-deficiency fails to confer atheroprotection in 52-week-old *Apoe^−/–^* mice. Plaque area was quantified in brachiocephalic artery (BCA), aortic root, aortic arch, thoracic and abdominal aorta of in 52-week-old *Apoe^−/–^Mif ^−/–^* mice that had been on HFD for the last 6 weeks before sacrifice (blue) and compared to corresponding Apoe–/–controls (grey). A-B) Representative images (A) and quantification (B) of ORO-stained sections of the aortic root. For each mouse, 12 serial sections with a distance of 50 µm were used for analysis. The mean plaque area is depicted in mm2 (n=3-5 mice per group; results are presented as means ± SD; scale bar: 200 µm). C-D) Representative images (C) and plaque quantification (D) of H&E-stained sections of the BCA. Serial sections were obtained as above. The mean plaque area is depicted as percentage of the total inner vessel area including the plaque (n=6- 7 mice per group; results are presented as means ± SD; scale bar: 100 µm). E) Representative images of en face-prepared ORO-stained aortas. F-H) Quantification of the plaque area in the aorta including aortic arch (F), thoracic aorta (G), and abdominal aorta (H). The plaque area is depicted as percentage of the total aortic surface (n=6-7 mice per group; results are presented as means ± SD). I-J) FACS-based quantification of total CD3e+ T cells (left panels), CD4+ T- cell subsets (middle panels), and CD8+ T-cell subsets (right panels) in spleen (I) and blood (J) of Apoe–/–Mif–/– mice (blue) and comparison to Apoe–/– controls (grey). 52-week-old mice on HFD for 6 weeks before sacrifice were examined (n=4-12 mice per group; results are presented as means ± SD). Statistics: Mann-Whitney Test, *, P<0.05, **, P<0.01, non- significant results with P values between 0.05 and 0.1 are indicated; each data point represents one mouse.

**Supplementary Figure S2:**
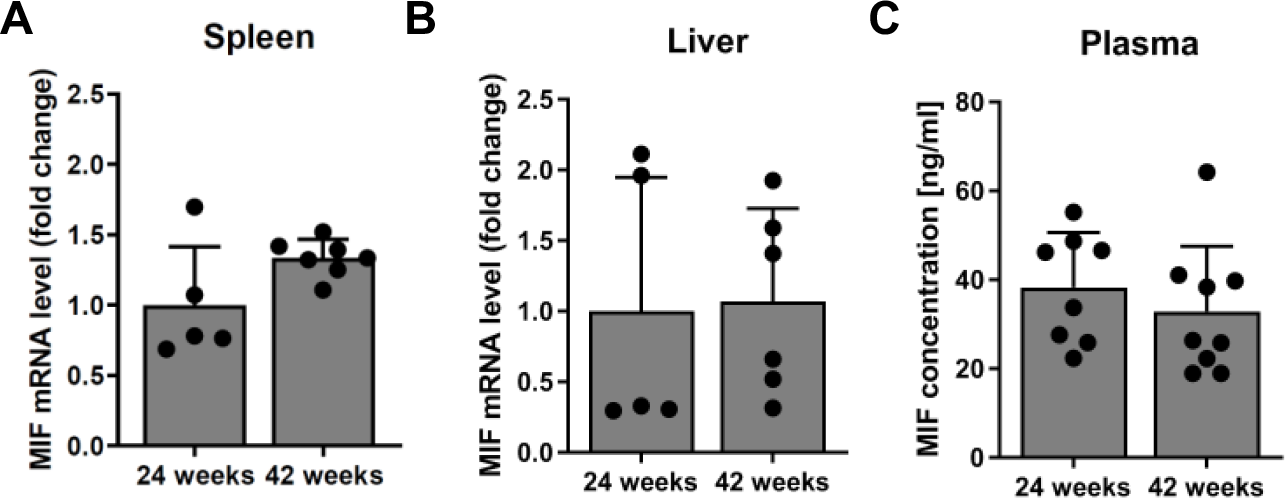
MIF levels remain unchanged with increased age and duration of HFD in *Apoe^−/–^* mice. **A-B**) mRNA expression of MIF was assessed by RT-qPCR in spleen (**A**) and liver (**B**) of *Apoe^−/–^* mice after 24 and 42 weeks of HFD. Values were normalized to the values of the 30-week-old *Apoe^−/–^* controls that received HFD for 24 weeks (n=5-7 mice; results are presented as means ± SD). **C**) MIF plasma levels as measured by ELISA in *Apoe^−/–^* mice after 24 and 42 weeks of HFD. Values were normalized to the *Apoe^−/–^* controls that received HFD for 24 weeks (n=8-9 mice per group; results are presented as means ± SD). Statistics: Mann-Whitney test (no significant differences observed; each data points represents one mouse).

**Supplementary Figure S3:**
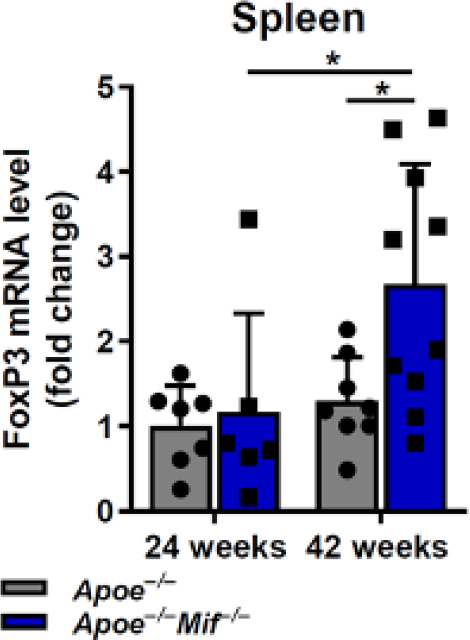
Increased numbers of splenic FoxP3^+^ Tregs in aged *Mif*- deficient *Apoe^−/–^* mice. mRNA expression of FoxP3 was assessed by RT-qPCR in spleen of *Apoe^−/–^Mif^−/–^* (blue) after 24 and 42 weeks of HFD and compared to the WT controls (grey). Values were normalized to the values of the 30-week-old *Apoe^−/–^* controls that received HFD for 24 weeks (n=6-10 mice per group; results are presented as means ± SD). Statistics: two- way ANOVA (each data point represents one mouse).

**Figure S4:**
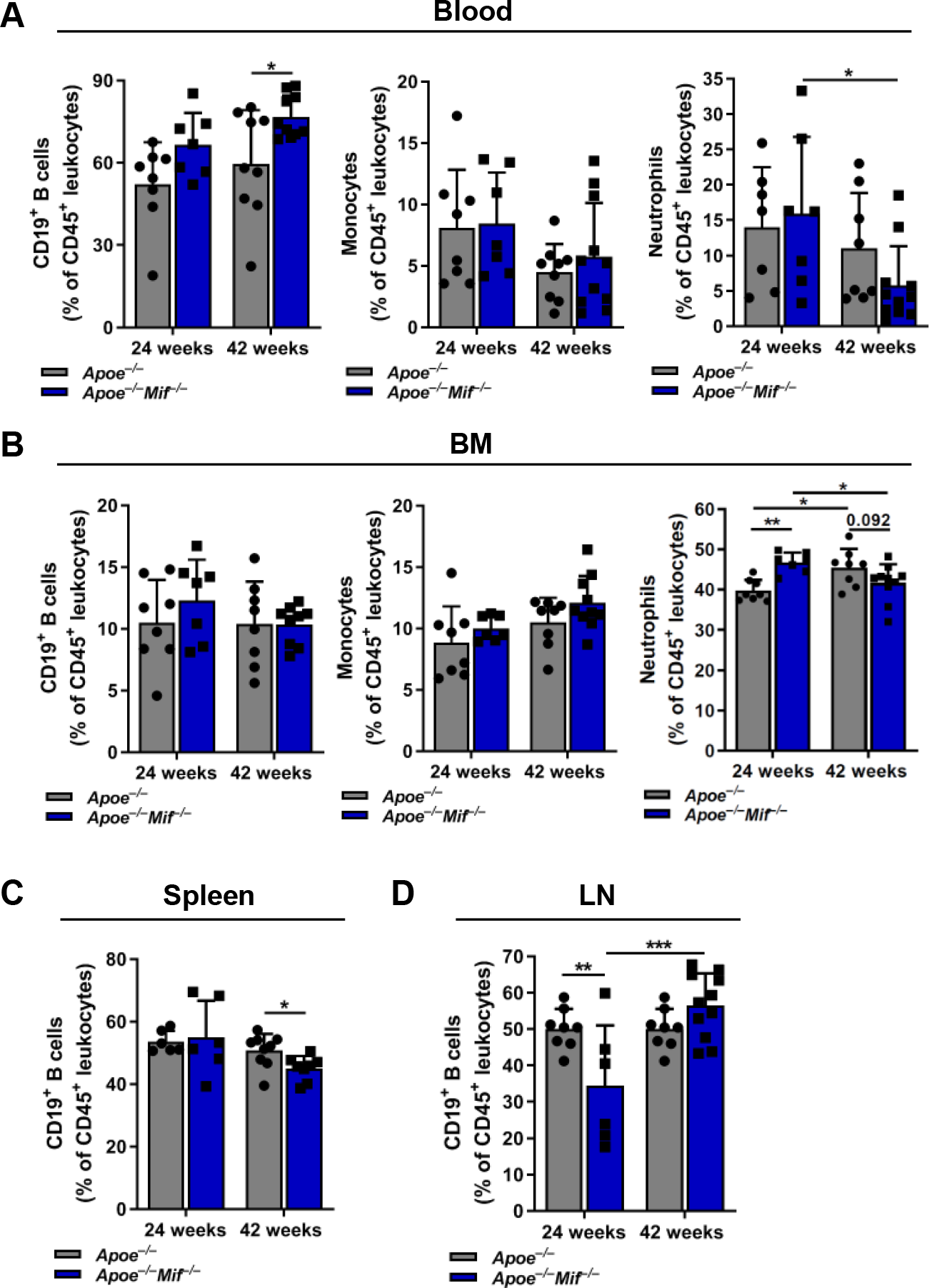
FACS-based analysis of immune cell numbers in immune organs and periphery. **A-D**) B cells (*left panels*), monocytes (*middle panels*) and neutrophils (*right panels*) were quantified by flow cytometry in the blood (**A**), BM (**D**), spleen **(C**) and LNs (**D**) of *Apoe^−/–^ Mif^−/–^* mice (blue) after 24 and 42 weeks of HFD and compared to the *Apoe^−/–^* controls. Monocyte and neutrophil counts in the spleen and LN were below 5 % and are not shown (n=6- 11, results are presented as means ± SD). Statistics: two-way ANOVA; *, *P* ≤ 0.05, **, *P* ≤ 0.01, ***, *P* ≤ 0.001, non-significant results with *P* values between 0.05 and 0.1 are given as precise numbers; each data point represents one independent mouse.

**Figure S5:**
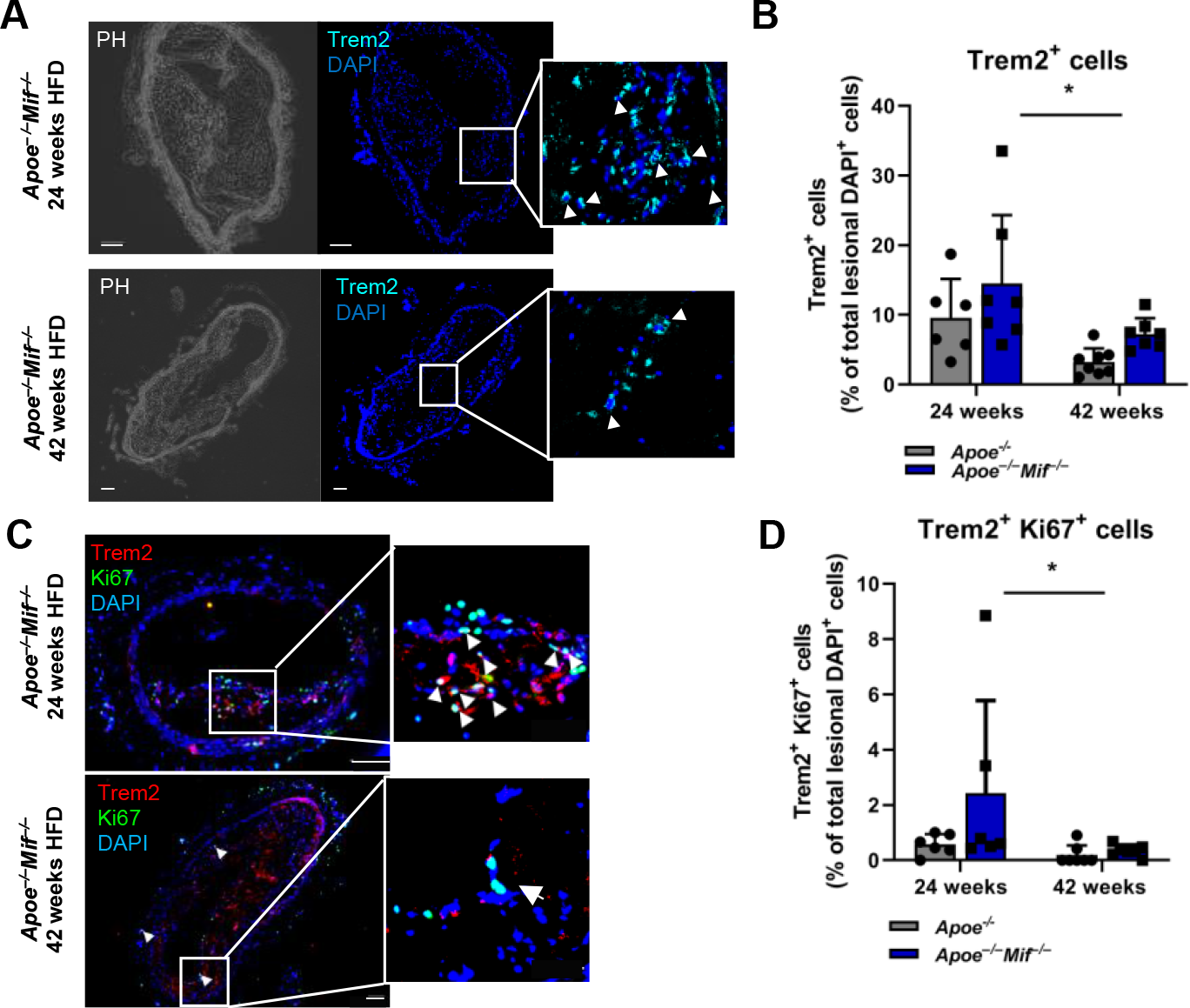
Age- and MIF-dependent changes in lesional Trem2^+^ macrophage numbers. **A-D**) Representative images (**A**) and quantification (**B**) of Trem2^+^ cells within the plaque. Cells are indicated by an arrow in the enlarged image section. Trem2^+^ cells were quantified manually and are depicted as percentage of total DAPI^+^ cells in the plaque. As Trem2 is mainly expressed by myeloid cells, the stained cells were considered as Trem2^+^ macrophages. **C-D)** Representative images (**C**) and quantification (**D**) of lesional Trem2^+^ Ki67^+^ cells. Trem2^+^ Ki67^+^ cells are indicated by arrows in the enlarged image section. Trem2^+^ Ki67^+^ double-positive cells were manually quantified and are depicted as percentage of total DAPI^+^ cells in the plaque. Results are presented as means ± SD from n=6-7 mice; scale bar: 100 µm. Statistics: two-way ANOVA; *, *P*<0.05; each data point represents one independent mouse. The data are identical to those in Figure 3E and G of the main manuscript, except that the quantification of Trem2^+^ and Trem2^+^ Ki67^+^ cells is expressed in relation to DAPI^+^ cells in the plaque area.

## References

1. Virani, S. S., Alonso, A., Benjamin, E. J., Bittencourt, M. S., Callaway, C. W., Carson, A. P., Chamberlain, A. M., Chang, A. R., Cheng, S., Delling, F. N., Djousse, L., Elkind, M. S. V., Ferguson, J. F., Fornage, M., Khan, S. S., Kissela, B. M., Knutson, K. L., Kwan, T. W., Lackland, D. T., Lewis, T. T., Lichtman, J. H., Longenecker, C. T., Loop, M. S., Lutsey, P. L., Martin, S. S., Matsushita, K., Moran, A. E., Mussolino, M. E., Perak, A. M., Rosamond, W. D., Roth, G. A., Sampson, U. K. A., Satou, G. M., Schroeder, E. B., Shah, S. H., Shay, C. M., Spartano, N. L., Stokes, A., Tirschwell, D. L., VanWagner, L. B., Tsao, C. W., American Heart Association Council on, E., Prevention Statistics, C., and Stroke Statistics, S. (2020) Heart disease and stroke statistics-2020 update: A report from the American Heart Association. Circulation 141, e139–e596

2. Joseph, P., Leong, D., McKee, M., Anand, S. S., Schwalm, J. D., Teo, K., Mente, A., and Yusuf, S. (2017) Reducing the global burden of cardiovascular disease, part 1: The epidemiology and risk factors. Circ Res 121, 677–694

3. Noels, H., Weber, C., and Koenen, R. R. (2019) Chemokines as therapeutic targets in cardiovascular disease. Arterioscler Thromb Vasc Biol 39, 583–592

4. Weber, C., and Noels, H. (2011) Atherosclerosis: current pathogenesis and therapeutic options. Nat Med 17, 1410–1422

5. Hansson, G. K., and Hermansson, A. (2011) The immune system in atherosclerosis. Nat Immunol 12, 204–212

6. Ross, R. (1999) Atherosclerosis--an inflammatory disease. N Engl J Med 340, 115–126

7. Kapurniotu, A., Gokce, O., and Bernhagen, J. (2019) The multitasking potential of alarmins and atypical chemokines. Front Med (Lausanne) 6, 3

8. Burger-Kentischer, A., Goebel, H., Seiler, R., Fraedrich, G., Schaefer, H. E., Dimmeler, S., Kleemann, R., Bernhagen, J., and Ihling, C. (2002) Expression of macrophage migration inhibitory factor in different stages of human atherosclerosis. Circulation 105, 1561–1566

9. Sinitski, D., Kontos, C., Krammer, C., Asare, Y., Kapurniotu, A., and Bernhagen, J. (2019) Macrophage migration inhibitory factor (MIF)-based therapeutic concepts in atherosclerosis and inflammation. Thromb Haemost 119, 553–566

10. Zernecke, A., Bernhagen, J., and Weber, C. (2008) Macrophage migration inhibitory factor in cardiovascular disease. Circulation 117, 1594–1602

11. Müller, I. I., Müller, K. A. L., Schönleber, H., Karathanos, A., Schneider, M., Jorbenadze, R., Bigalke, B., Gawaz, M., and Geisler, T. (2012) Macrophage migration inhibitory factor is enhanced in acute coronary syndromes and is associated with the inflammatory response. PLoS One 7, e38376

12. Calandra, T., and Roger, T. (2003) Macrophage migration inhibitory factor: a regulator of innate immunity. Nat Rev Immunol 3, 791–800

13. Tillmann, S., Bernhagen, J., and Noels, H. (2013) Arrest functions of the MIF ligand/receptor axes in atherogenesis. Front Immunol 4, 115

14. Adamali, H., Armstrong, M. E., McLaughlin, A. M., Cooke, G., McKone, E., Costello, C. M., Gallagher, C. G., Leng, L., Baugh, J. A., Fingerle-Rowson, G., Bucala, R. J., McLoughlin, P., and Donnelly, S. C. (2012) Macrophage migration inhibitory factor enzymatic activity, lung inflammation, and cystic fibrosis. Am J Respir Crit Care Med 186, 162–169

15. Baugh, J. A., Chitnis, S., Donnelly, S. C., Monteiro, J., Lin, X., Plant, B. J., Wolfe, F., Gregersen, P. K., and Bucala, R. (2002) A functional promoter polymorphism in the macrophage migration inhibitory factor (MIF) gene associated with disease severity in rheumatoid arthritis. Genes Immun 3, 170–176

16. Lefebvre, J. S., Maue, A. C., Eaton, S. M., Lanthier, P. A., Tighe, M., and Haynes, L. (2012) The aged microenvironment contributes to the age-related functional defects of CD4 T cells in mice. Aging Cell 11, 732–740

17. Wirtz, T. H., Tillmann, S., Strussmann, T., Kraemer, S., Heemskerk, J. W., Grottke, O., Gawaz, M., von Hundelshausen, P., and Bernhagen, J. (2015) Platelet-derived MIF: a novel platelet chemokine with distinct recruitment properties. Atherosclerosis 239, 1–10

18. Bernhagen, J., Krohn, R., Lue, H., Gregory, J. L., Zernecke, A., Koenen, R. R., Dewor, M., Georgiev, I., Schober, A., Leng, L., Kooistra, T., Fingerle-Rowson, G., Ghezzi, P., Kleemann, R., McColl, S. R., Bucala, R., Hickey, M. J., and Weber, C. (2007) MIF is a noncognate ligand of CXC chemokine receptors in inflammatory and atherogenic cell recruitment. Nat Med 13, 587–596

19. Asare, Y., Schmitt, M., and Bernhagen, J. (2013) The vascular biology of macrophage migration inhibitory factor (MIF). Expression and effects in inflammation, atherogenesis and angiogenesis. Thromb Haemost 109, 391–398

20. Schober, A., Bernhagen, J., Thiele, M., Zeiffer, U., Knarren, S., Roller, M., Bucala, R., and Weber, C. (2004) Stabilization of atherosclerotic plaques by blockade of macrophage migration inhibitory factor after vascular injury in apolipoprotein E-deficient mice. Circulation 109, 380–385

21. A. Pan, J. H., Sukhova, G. K., Yang, J. T., Wang, B., Xie, T., Fu, H., Zhang, Y., Satoskar, A. R., David, J. R., Metz, C. N., Bucala, R., Fang, K., Simon, D. I., Chapman, H. A., Libby, P., and Shi, G. P. (2004) Macrophage migration inhibitory factor deficiency impairs atherosclerosis in low-density lipoprotein receptor-deficient mice. Circulation 109, 3149–3153

22. Burger-Kentischer, A., Gobel, H., Kleemann, R., Zernecke, A., Bucala, R., Leng, L., Finkelmeier, D., Geiger, G., Schaefer, H. E., Schober, A., Weber, C., Brunner, H., Rutten, H., Ihling, C., and Bernhagen, J. (2006) Reduction of the aortic inflammatory response in spontaneous atherosclerosis by blockade of macrophage migration inhibitory factor (MIF). Atherosclerosis 184, 28–38

23. Schmitz, C., Noels, H., El Bounkari, O., Straussfeld, E., Megens, R. T. A., Sternkopf, M., Alampour-Rajabi, S., Krammer, C., Tilstam, P. V., Gerdes, N., Burger, C., Kapurniotu, A., Bucala, R., Jankowski, J., Weber, C., and Bernhagen, J. (2018) Mif-deficiency favors an atheroprotective autoantibody phenotype in atherosclerosis. FASEB J 32, 4428–4443

24. Head, T., Daunert, S., and Goldschmidt-Clermont, P. J. (2017) The Aging Risk and Atherosclerosis: A Fresh Look at Arterial Homeostasis. Front Genet 8, 216

25. Liberale, L., Montecucco, F., Tardif, J. C., Libby, P., and Camici, G. G. (2020) Inflamm- ageing: the role of inflammation in age-dependent cardiovascular disease. Eur Heart J 41, 2974–2982

26. Nakashima, Y., Plump, A. S., Raines, E. W., Breslow, J. L., and Ross, R. (1994) ApoE-deficient mice develop lesions of all phases of atherosclerosis throughout the arterial tree. Arterioscler Thromb Vasc Biol 14, 133–140

27. Fox, J., Barthold, S., Davisson, M., Newcomer, C., Quimby, F., and Smith, A. (2007) The Mouse in Biomedical Research, Elsevier, ISBN: 9780080469089

28. Mohanta, S. K., Yin, C., Peng, L., Srikakulapu, P., Bontha, V., Hu, D., Weih, F., Weber, C., Gerdes, N., and Habenicht, A. J. (2014) Artery tertiary lymphoid organs contribute to innate and adaptive immune responses in advanced mouse atherosclerosis. Circ Res 114, 1772–1787

29. Yin, C., Mohanta, S. K., Srikakulapu, P., Weber, C., and Habenicht, A. J. (2016) Artery tertiary lymphoid organs: powerhouses of atherosclerosis immunity. Front Immunol 7, 387

30. Srikakulapu, P., Hu, D., Yin, C., Mohanta, S. K., Bontha, S. V., Peng, L., Beer, M., Weber, C., McNamara, C. A., Grassia, G., Maffia, P., Manz, R. A., and Habenicht, A.J. (2016) Artery tertiary lymphoid organs control multilayered territorialized atherosclerosis B-cell responses in aged ApoE-/- mice. Arterioscler Thromb Vasc Biol 36, 1174–1185

31. Akhavanpoor, M., Gleissner, C. A., Akhavanpoor, H., Lasitschka, F., Doesch, A. O., Katus, H. A., and Erbel, C. (2018) Adventitial tertiary lymphoid organ classification in human atherosclerosis. Cardiovasc Pathol 32, 8–14

32. Fingerle-Rowson, G., Petrenko, O., Metz, C. N., Forsthuber, T. G., Mitchell, R., Huss, R., Moll, U., Muller, W., and Bucala, R. (2003) The p53-dependent effects of macrophage migration inhibitory factor revealed by gene targeting. Proc Natl Acad Sci U S A 100, 9354–9359

33. Chen, Z., Sakuma, M., Zago, A. C., Zhang, X., Shi, C., Leng, L., Mizue, Y., Bucala, R., and Simon, D. (2004) Evidence for a role of macrophage migration inhibitory factor in vascular disease. Arterioscler Thromb Vasc Biol 24, 709–714

34. Ma, H., Wang, J., Thomas, D. P., Tong, C., Leng, L., Wang, W., Merk, M., Zierow, S., Bernhagen, J., Ren, J., Bucala, R., and Li, J. (2010) Impaired macrophage migration inhibitory factor-AMP-activated protein kinase activation and ischemic recovery in the senescent heart. Circulation 122, 282–292

35. Cochain, C., Vafadarnejad, E., Arampatzi, P., Pelisek, J., Winkels, H., Ley, K., Wolf, D., Saliba, A. E., and Zernecke, A. (2018) Single-cell RNA-seq reveals the transcriptional landscape and heterogeneity of aortic macrophages in murine atherosclerosis. Circ Res 122, 1661–1674

36. Willemsen, L., and de Winther, M. P. (2020) Macrophage subsets in atherosclerosis as defined by single-cell technologies. J Pathol 250, 705–714

37. Kim, K., Shim, D., Lee, J. S., Zaitsev, K., Williams, J. W., Kim, K. W., Jang, M. Y., Seok Jang, H., Yun, T. J., Lee, S. H., Yoon, W. K., Prat, A., Seidah, N. G., Choi, J., Lee, S. P., Yoon, S. H., Nam, J. W., Seong, J. K., Oh, G. T., Randolph, G. J., Artyomov, M. N., Cheong, C., and Choi, J. H. (2018) Transcriptome analysis reveals nonfoamy rather than foamy plaque macrophages are proinflammatory in atherosclerotic murine models. Circ Res 123, 1127–1142

38. Campbell, K. A., Lipinski, M. J., Doran, A. C., Skaflen, M. D., Fuster, V., and McNamara, C. A. (2012) Lymphocytes and the adventitial immune response in atherosclerosis. Circ Res 110, 889–900

39. Hu, D., Mohanta, S. K., Yin, C., Peng, L., Ma, Z., Srikakulapu, P., Grassia, G., MacRitchie, N., Dever, G., Gordon, P., Burton, F. L., Ialenti, A., Sabir, S. R., McInnes, I. B., Brewer, J. M., Garside, P., Weber, C., Lehmann, T., Teupser, D., Habenicht, L., Beer, M., Grabner, R., Maffia, P., Weih, F., and Habenicht, A. J. (2015) Artery tertiary lymphoid organs control aorta immunity and protect against atherosclerosis via vascular smooth muscle cell lymphotoxin beta receptors. Immunity 42, 1100–1115

40. Tsiantoulas, D., Diehl, C. J., Witztum, J. L., and Binder, C. J. (2014) B cells and humoral immunity in atherosclerosis. Circ Res 114, 1743–1756

41. Tyrrell, D. J., and Goldstein, D. R. (2021) Ageing and atherosclerosis: vascular intrinsic and extrinsic factors and potential role of IL-6. Nat Rev Cardiol 18, 58–68

42. Liu, D., Richardson, G., Benli, F. M., Park, C., de Souza, J. V., Bronowska, A. K., and Spyridopoulos, I. (2020) Inflammageing in the cardiovascular system: mechanisms, emerging targets, and novel therapeutic strategies. Clin Sci (Lond) 134, 2243–2262

43. Smykiewicz, P., Segiet, A., Keag, M., and Zera, T. (2018) Proinflammatory cytokines and ageing of the cardiovascular-renal system. Mech Ageing Dev 175, 35–45

44. Rammos, C., Hendgen-Cotta, U. B., Pohl, J., Totzeck, M., Luedike, P., Schulze, V. T., Kelm, M., and Rassaf, T. (2014) Modulation of circulating macrophage migration inhibitory factor in the elderly. Biomed Res Int 2014, 582586

45. Rammos, C., Hendgen-Cotta, U. B., Deenen, R., Pohl, J., Stock, P., Hinzmann, C., Kelm, M., and Rassaf, T. (2014) Age-related vascular gene expression profiling in mice. Mech Ageing Dev 135, 15–23

46. A. Trott, D. W., Henson, G. D., Ho, M. H. T., Allison, S. A., Lesniewski, L. A., and Donato, A. J. (2018) Age-related arterial immune cell infiltration in mice is attenuated by caloric restriction or voluntary exercise. Exp Gerontol 109, 99–107

47. Trott, D. W., and Fadel, P. J. (2019) Inflammation as a mediator of arterial ageing. Exp Physiol 104, 1455–1471

48. Du, W., Wong, C., Song, Y., Shen, H., Mori, D., Rotllan, N., Price, N., Dobrian, A. D., Meng, H., Kleinstein, S. H., Fernandez-Hernando, C., and Goldstein, D. R. (2016) Age-associated vascular inflammation promotes monocytosis during atherogenesis. Aging Cell 15, 766–777

49. Castro, B. A., Flanigan, P., Jahangiri, A., Hoffman, D., Chen, W., Kuang, R., De Lay, M., Yagnik, G., Wagner, J. R., Mascharak, S., Sidorov, M., Shrivastav, S., Kohanbash, G., Okada, H., and Aghi, M. K. (2017) Macrophage migration inhibitory factor downregulation: a novel mechanism of resistance to anti-angiogenic therapy. Oncogene 36, 3749–3759

50. Kim, B. S., Tilstam, P. V., Arnke, K., Leng, L., Ruhl, T., Piecychna, M., Schulte, W., Sauler, M., Frueh, F. S., Storti, G., Lindenblatt, N., Giovanoli, P., Pallua, N., Bernhagen, J., and Bucala, R. (2020) Differential regulation of macrophage activation by the MIF cytokine superfamily members MIF and MIF-2 in adipose tissue during endotoxemia. FASEB J 34, 4219–4233

51. Aw, D., Hilliard, L., Nishikawa, Y., Cadman, E. T., Lawrence, R. A., and Palmer, D. B. (2016) Disorganization of the splenic microanatomy in ageing mice. Immunology 148, 92–101

52. Masters, A. R., Jellison, E. R., Puddington, L., Khanna, K. M., and Haynes, L. (2018) Attrition of T Cell Zone Fibroblastic Reticular Cell Number and Function in Aged Spleens. ImmunoHorizons 2, 155–163

53. Moos, M. P., John, N., Grabner, R., Nossmann, S., Gunther, B., Vollandt, R., Funk, C. D., Kaiser, B., and Habenicht, A. J. (2005) The lamina adventitia is the major site of immune cell accumulation in standard chow-fed apolipoprotein E-deficient mice. Arterioscler Thromb Vasc Biol 25, 2386–2391

54. Stranford, S., and Ruddle, N. H. (2012) Follicular dendritic cells, conduits, lymphatic vessels, and high endothelial venules in tertiary lymphoid organs: Parallels with lymph node stroma. Front Immunol 3, 350

55. Kyaw, T., Tay, C., Krishnamurthi, S., Kanellakis, P., Agrotis, A., Tipping, P., Bobik, A., and Toh, B. H. (2011) B1a B lymphocytes are atheroprotective by secreting natural IgM that increases IgM deposits and reduces necrotic cores in atherosclerotic lesions. Circ Res 109, 830–840

56. Rosenfeld, S. M., Perry, H. M., Gonen, A., Prohaska, T. A., Srikakulapu, P., Grewal, S., Das, D., McSkimming, C., Taylor, A. M., Tsimikas, S., Bender, T. P., Witztum, J. L., and McNamara, C. A. (2015) B-1b cells secrete atheroprotective IgM and attenuate atherosclerosis. Circ Res 117, e28–39

57. Ridker, P. M., Everett, B. M., Thuren, T., MacFadyen, J. G., Chang, W. H., Ballantyne, C., Fonseca, F., Nicolau, J., Koenig, W., Anker, S. D., Kastelein, J. J. P., Cornel, J. H., Pais, P., Pella, D., Genest, J., Cifkova, R., Lorenzatti, A., Forster, T., Kobalava, Z., Vida-Simiti, L., Flather, M., Shimokawa, H., Ogawa, H., Dellborg, M., Rossi, P. R. F., Troquay, R. P. T., Libby, P., Glynn, R. J., and Group, C. T. (2017) Antiinflammatory therapy with canakinumab for atherosclerotic disease. N Engl J Med 377, 1119–1131

58. Kontos, C., El Bounkari, O., Krammer, C., Sinitski, D., Hille, K., Zan, C., Yan, G., Wang, S., Gao, Y., Brandhofer, M., Megens, R. T. A., Hoffmann, A., Pauli, J., Asare, Y., Gerra, S., Bourilhon, P., Leng, L., Eckstein, H. H., Kempf, W. E., Pelisek, J., Gokce, O., Maegdefessel, L., Bucala, R., Dichgans, M., Weber, C., Kapurniotu, A., and Bernhagen, J. (2020) Designed CXCR4 mimic acts as a soluble chemokine receptor that blocks atherogenic inflammation by agonist-specific targeting. Nat Commun 11, 5981

59. Krammer, C., Kontos, C., Dewor, M., Hille, K., Dalla Volta, B., El Bounkari, O., Tas, K., Sinitski, D., Brandhofer, M., Megens, R. T. A., Weber, C., Schultz, J. R., Bernhagen, J., and Kapurniotu, A. (2020) A MIF-derived cyclopeptide that inhibits MIF binding and atherogenic signaling via the chemokine receptor CXCR2. Chembiochem 22, 1012–1019

60. Merk, M., Zierow, S., Leng, L., Das, R., Du, X., Schulte, W., Fan, J., Lue, H., Chen, Y., Xiong, H., Chagnon, F., Bernhagen, J., Lolis, E., Mor, G., Lesur, O., and Bucala, R. (2011) The D-dopachrome tautomerase (DDT) gene product is a cytokine and functional homolog of macrophage migration inhibitory factor (MIF). Proc Natl Acad Sci U S A 108, E577–585

61. Merk, M., Mitchell, R. A., Endres, S., and Bucala, R. (2012) D-dopachrome tautomerase (D-DT or MIF-2): Doubling the MIF cytokine family. Cytokine 59, 10–17

62. Averdunk, L., Bernhagen, J., Fehnle, K., Surowy, H., Ludecke, H. J., Mucha, S., Meybohm, P., Wieczorek, D., Leng, L., Marx, G., Leaf, D. E., Zarbock, A., Zacharowski, K., On Behalf Of The, R. S. C., Bucala, R., and Stoppe, C. (2020) The macrophage migration inhibitory factor (MIF) promoter polymorphisms (rs3063368, rs755622) predict acute kidney injury and death after cardiac surgery. J Clin Med 9, 2936

63. Lan, M. Y., Chang, Y. Y., Chen, W. H., Tseng, Y. L., Lin, H. S., Lai, S. L., and Liu, J. S. (2013) Association between MIF gene polymorphisms and carotid artery atherosclerosis. Biochem Biophys Res Commun 435, 319–322

